# YAP promotes chromosomal instability by transcriptionally activating AJUBA through its super enhancer in breast cancer

**DOI:** 10.1101/2024.02.19.579918

**Authors:** Qingwen Huang, Rui Zhang, Weijian Meng, Hongliang Dong, Zhihong Qi, Zhuo Chen, Liang Liu, Jie Shen, Daxing Xie

**Author notes:** Correspondence: Jie Shen and Daxing Xie, Tongji Hospital, Tongji Medical College, Huazhong University of Science and Technology, 1095 Jiefang Ave. Wuhan, Hubei 430030, China. and. These authors contribute equally to this work (Qingwen Huang, Rui Zhang).

## Abstract

Chromosomal instability (CIN) has been proven to be a fundamental property of cancer and a key underlying mechanism of tumorigenesis and progression. Still, relatively few studies have revealed the mechanism of this process. In this research, we identified the core Hippo component, YAP, played a crucial role in CIN formation in breast cancer. Using public ChIP-seq data and bioinformatic analysis, we found YAP could transcriptionally induce spindle assembly checkpoint, AJUBA, via TEAD, then activate AJUBA-AURKA signaling. Abolish AJUBA-AURKA axis could reverse YAP-induced CIN formation and aberrant mitosis. Mechanically, YAP/TEAD complex could accumulate to the super enhancer region of AJUBA, thus significantly induced its expression, and subsequently promoted AURKA auto-phosphorylating. These findings revealed a novel cross-talk between Hippo and AJUBA-AURKA signaling, and provided a new insight in CIN formation in breast cancer.

## Introduction

Chromosomal instability is a common form of genomic instability which presents as numerical or structural alterations of chromosome, and leads to loss/amplification of driver genes, focal rearrangement, extrachromosomal DNA, micronuclei formation and innate immune activation in various cancer types^1^. Therefore, CIN is tightly associated with tumorigenesis, tumor growth, metastasis, chemotherapy resistance and poor prognosis^2^. The causes of CIN are diverse and complex, including mitotic errors (dysfunction of kinetochore-microtubule attachment; dysregulation of sister chromatids cohesion and supernumerary centrosomes), replication stress (cell cycle regulator dysfunction), homologous recombination deficiency, telomere crisis and repeated cycles of chromosome breakage/rearrangement (spindle assembly checkpoint gene mutation)^3–7^. Due to the diversity of biological processes involved in CIN, the specific oncogenic signaling pathways which trigger CIN occurrence are still uncertain.

Recently, Hippo/YAP signaling is reported to participate in CIN regulation^8^. YAP, as well as its paralogue TAZ, are often acted as a nuclear transcriptional coactivator to regulate organ growth and tissue homeostasis, and are closely related to tumor malignancy and poor prognosis in multiple cancers^9–12^. In previous researches, YAP was observed to be associated with CIN in cervical cancer and cholangiocarcinoma^8,13^. Moreover, YAP was found to induce CIN-related gene expression, centrosomes abnormal duplication and polyploid formation^14–16^. Current viewpoint is that YAP could induce Forkhead transcription factor, FOXM1, expression through its promoter and trigger aneuploid phenotype^14^. However, the further specific mechanism of YAP regulated CIN remains to be explored.

According to conventional viewpoints, YAP/TAZ often partners nuclear transcription factors (e.g. TEADs) to interacts with the promoter region of downstream target gene and induces its expression^17^. In 2015, Zanconato F et al. reported that YAP-TEADs complex could synergistically regulate target genes which was directly involved in S-phase entry and mitosis, however, this control occurred almost exclusively from enhancers that activated target promoters through chromatin looping^18^. Further research also reported that YAP/TAZ-bound enhancers acted as key for the cancer cell state and presented active chromatin profiles across diverse human tumors^19^. Some enhancers were highly enriched in a continuous region of genome, and have a strong ability to regulate cell state and biological functions, thus these gene loci were named as “super enhancers (SEs)”^20^. YAP/TAZ was observed to regulate downstream biological process via modulating SEs in various cell type^21–23^. These findings indicate that enhancers, especially super enhancer, might play essential role in YAP signaling. In this study, we found YAP/TAZ-TEADs could transcriptionally induce spindle assembly checkpoint, AJUBA, expression, and activate AJUBA-AURKA signaling to directly induce abnormal mitosis and CIN formation in breast cancer. Mechanically, YAP/TEAD complex could accumulate to the super enhancer region of AJUBA via liquid-liquid phase separation, subsequently strengthens the AJUBA promoter-SE looping structure, thus induce its transcription and promoted AURKA auto-phosphorylating. These findings reveal a novel function and mechanism of YAP induced CIN in breast cancer, and presents a new insight in Hippo signaling related transcriptional activation through super enhancer system.

## Results

### YAP induces CIN formation and aberrant mitosis in breast cancer

To validate whether YAP correlated with CIN and aberrant mitosis, we first performed YAP IHC staining in a tissue array containing 75 primary breast cancer specimens and categorized these samples into YAP low expression group (n=35) and YAP high expression group (n=40) according to the IHC score, meanwhile, we examined and calculated the aberrant mitosis rates of these sample via oil immersion, and found tumor cells in YAP high expression group appeared a higher aberrant mitosis rate than in YAP low expression group (Fig. 1A).

**Figure 1.**
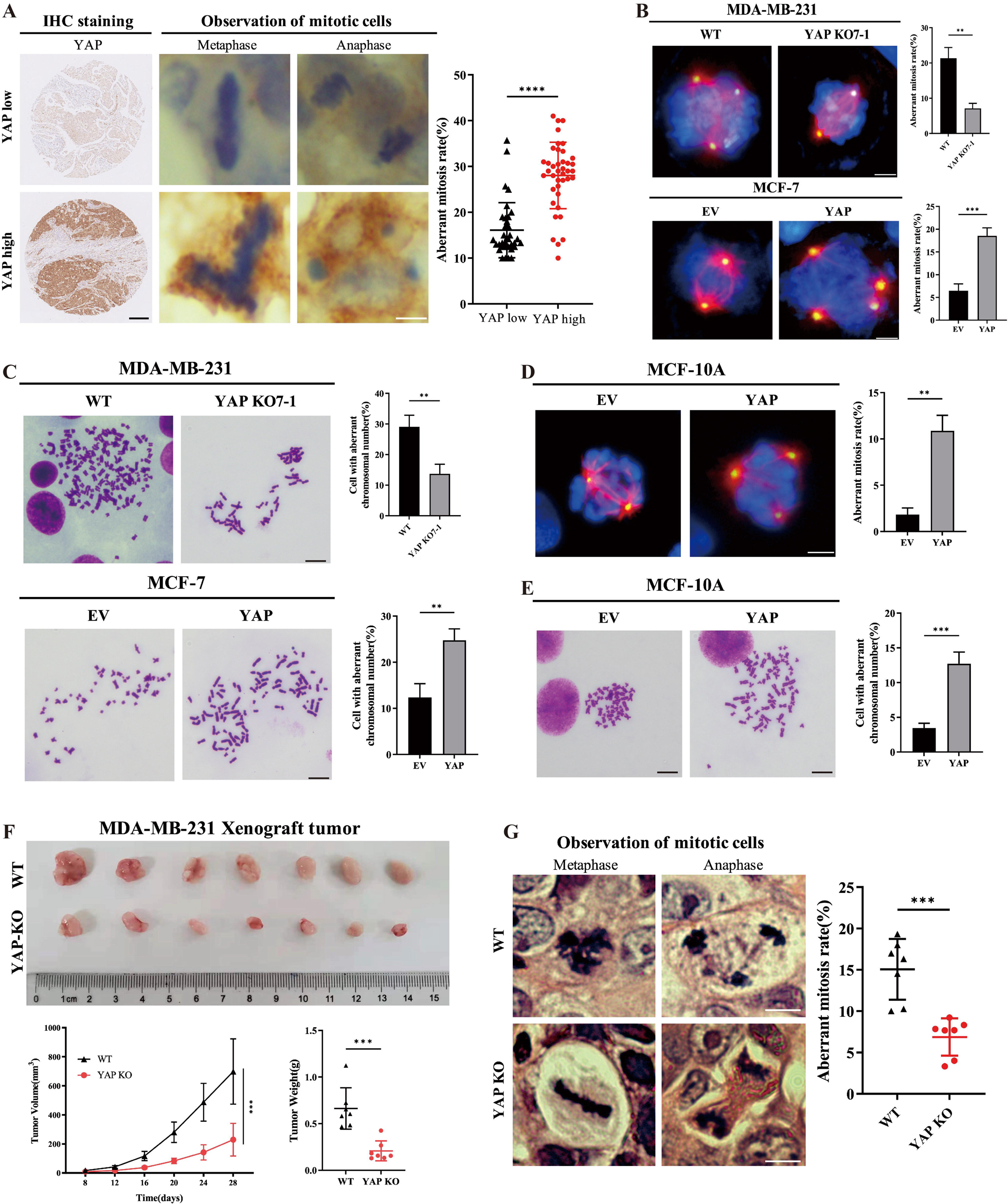
YAP induces CIN formation and aberrant mitosis in breast cancer. (A) Representative images of YAP immunohistochemistry staining (scale bar: 250μm) and aberrant mitotic cells (scale bar: 4μm) from human triple negative breast cancer tissue microarray (n=75). Scatterplot showed the mean percentage ± SD of aberrant mitosis rate. YAP low, YAP low expression group (YAP IHC score ≤ 5, n=35); YAP high, YAP high expression group (YAP IHC score > 6, n=40); ****p<0.001. (B) Aberrant mitosis was observed by immunofluorescence in MDA-MB-231 cells with wild type (WT) or YAP knocking out (YAP KO7-1) (upper); and MCF-7 cells stably transfected with empty vector (EV) or YAP overexpression plasmid (YAP) (lower). β-Tubulin were stained with Alexa Fluor 555 (red), γ-Tubulin were stained with Dylight 488 (green), and Nuclei were stained with DAPI (blue). Scale bar: 10μm. Histograms showed the mean percentage ± SD of aberrant mitosis rate. **p<0.01; ***p<0.001. (C) Representative images of chromosome metaphase spreading in MDA-MB-231 cells with wild type (WT) or YAP knocking out (YAP KO7-1) (upper); and MCF-7 cells stably transfected with empty vector (EV) or YAP overexpression plasmid (YAP) (lower). Scale bar: 5μm. Histograms showed the mean percentage ± SD of cell rate with aberrant chromosomal number. **p<0.01. (D) Aberrant mitosis in MCF-10A cells transfected with empty vector (EV) or YAP overexpression plasmid (YAP). β-Tubulin (red); γ-Tubulin (green); Nuclei (blue). Scale bar: 10μm. Histogram showed the mean percentage ± SD of aberrant mitosis rate. **p<0.01. (E) Chromosome metaphase spreading in MCF-10A cells transfected with empty vector (EV) or YAP overexpression plasmid (YAP). Scale bar: 5μm. Histogram shows the mean percentage ± SD of cell rate with aberrant chromosomal number. ***p<0.001. (F) BALB/c nude mice (n=7) were orthotopically implanted with MDA-MB-231 wild type cells (WT) or YAP knocking out cells (YAP-KO). YAP knocking out cells were the mixtures of two YAP-KO clones (YAP-KO-6-2 and YAP-KO-7-1). Tumor growth curve and tumor weights were measured as indicated. Data were presented as mean ± SD. ***p<0.001. (G) H&E staining of xenograft tumors from (H). Scale bar:4μm. Scatterplot showed the mean percentage ± SD of aberrant mitosis rate. ***p<0.001.

Next, we examined whether YAP could regulate CIN in vitro. As previously reported, YAP protein was relatively highly expressed in MDA-MB-231, and low show low expression in MCF7 and MCF10A^24^, thus these three cell lines were selected for further study. Firstly, we knocked out endogenous YAP via CRISPR/Cas9 in MDA-MB-231 cells and stable overexpressing YAP in MCF7 and MCF10A cells (Fig. S1A-C). Subsequently, paclitaxel was used to trigger mitotic arrest and the multipolar mitotic spindles was detected via immunofluorescence. Through microscopic observation, we found knocking out YAP could significantly reduce the proportion of cells with multipolar mitosis in MDA-MB-231 cell lines, in contrast, overexpressing YAP could significantly induce aberrant mitosis in MCF7cells (Fig. 1B). Through karyotype analysis, YAP expression level appeared to be positively correlated with aneuploid rate in both MDA-MB-231 and MCF7 cell lines (Fig. 1C). Similarly, overexpressing exogenous YAP could induce multipolar mitosis and aneuploid in non-tumorigenic breast epithelial cell line, MCF-10A (Fig. 1D, E). As homolog protein of YAP, TAZ presented a similar biological function in CIN formation. Therefore, we also knocked out endogenous TAZ via CRISPR/Cas9 in MDA-MB-231 cells and stable overexpressing TAZ in MCF7 and MCF10A cells (Fig. S1D-F). Immunofluorescence and karyotype analysis showed that knocking down TAZ could significantly decrease the proportion of cells with multipolar mitosis and aneuploid rate in MDA-MB-231 cells, while overexpressing TAZ could induce aberrant mitosis in MCF7 and MCF-10A cells (S.Fig. 1G-L)

To unveil the correlation between YAP and CIN in vivo, we established subcutaneous xenograft models of MDA-MB-231 on female NOD/SCID mice. Four weeks after tumor implantation, tumors were collected and analyzed. Tumor size measurement showed that knocking out YAP could significantly inhibit tumor growth in MDA-MB-231 (Fig.1F). Additionally, the proportion of aberrant mitosis was remarkably decreased in MDA-MB-231 xenografted tumor with YAP knockout (Fig.1G). These data demonstrated that YAP could induce aberrant mitosis and increase aneuploid rate, thus promote CIN formation in breast cancer cells.

### YAP-TEAD interaction is essential in CIN formation and aberrant mitosis

As a transcriptional co-activator, YAP/TAZ mostly interacted with transcriptional factor, TEADs, to enhance downstream genes transcription. To investigate whether TEADs involves in the YAP/TAZ induced CIN and aberrant mitosis. We first established MCF7 cells with stable exogenous YAP wild type and TEAD binding domain mutant (YAP-S94A) expression (Fig. S2A). YAP could significantly induce aberrant mitosis and aneuploidy formation, while this effect was abrogated in YAP-S94A group in MCF7 cells (Fig. 2A, B). In TEADs family, TEAD4 protein was dramatically transcribed and was positively correlated with poor prognosis in breast cancer^24^. Here, we transiently knock down TEAD4 in MCF7 with YAP/TAZ overexpression (Fig. S2B, C) or in MDA-MB-231 cells (Fig. S2D). The results showed that knocking down TEAD4 could significantly reverse YAP/TAZ induced aberrant mitosis and aneuploidy formation (Fig. 2C, D and Fig. S2E, F). Moreover, MCF7-YAP and MDA-MB-231 cells were treated with verteporfin (small molecular inhibitor of YAP/TAZ-TEADs interaction) and TED-347 (YAP/TAZ-TEAD4 specific dissociation agent) (Fig. 2SG, H), we found inhibition of YAP-TEADs interaction could also decrease the proportion of cells with multipolar mitosis and aneuploid rate (Fig. 2E-H). These results indicated that TEADs was essential in YAP/TAZ induced CIN in breast cancer.

**Figure 2.**
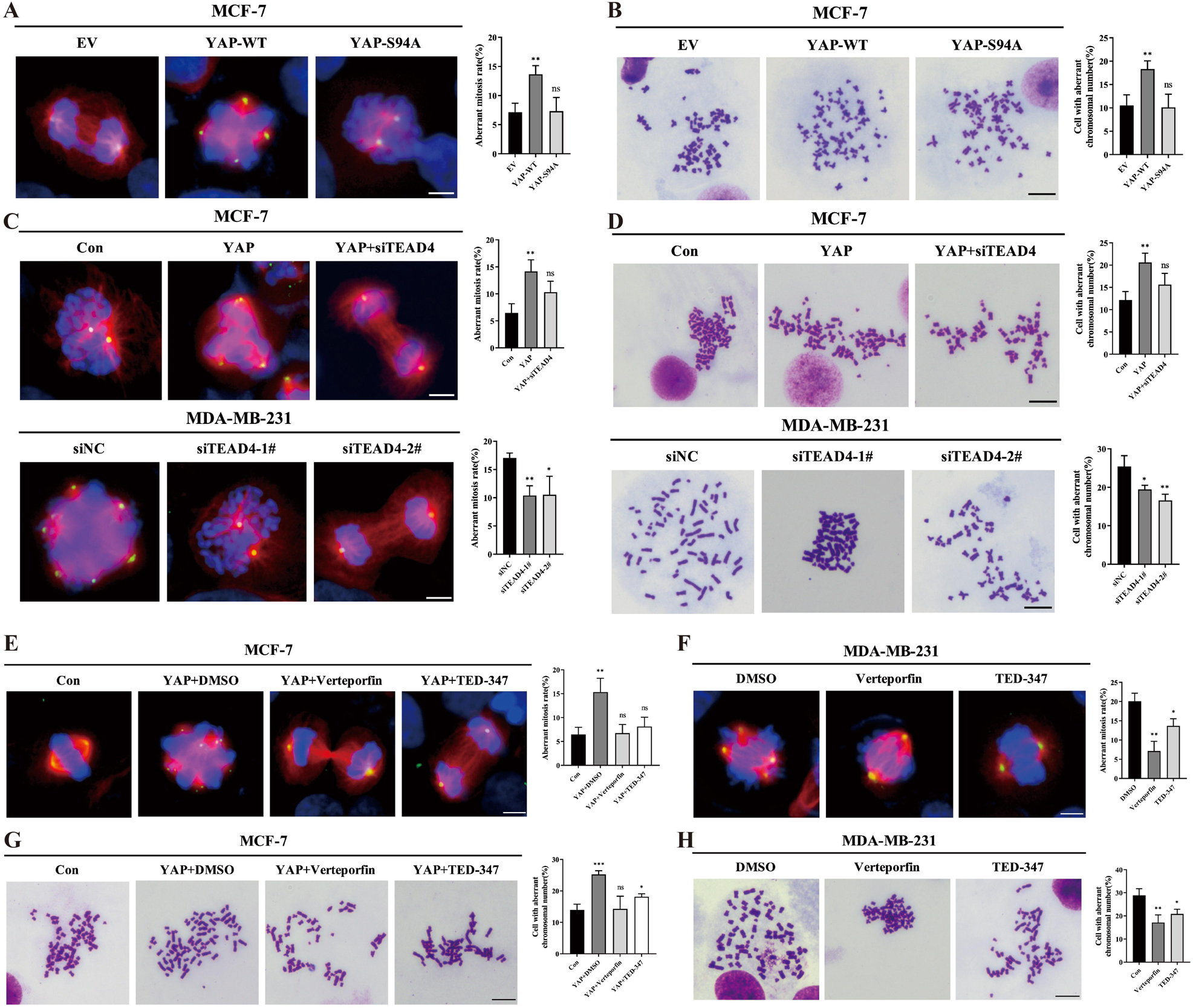
YAP-TEAD interaction is essential in CIN formation and aberrant mitosis. (A) Aberrant mitosis was observed by immunofluorescence in MCF7 cells transfected with empty vector (EV), wild type YAP (YAP-WT) or YAP-S94A mutant (YAP-S94A). β-Tubulin (red), γ-Tubulin (green), Nuclei (blue). Scale bar: 10μm. Histograms showed the mean percentage ± SD of aberrant mitosis rate. ns, no significant; **p<0.01; ***p<0.001. (B) Images of chromosome metaphase spreading in MCF7 cells transfected with empty vector (EV), wild type YAP (YAP-WT) or YAP-S94A mutant (YAP-S94A). Scale bar: 5μm. Histograms showed the mean percentage ± SD of cell rate with aberrant chromosomal number. ns, no significant; **p<0.01; ***p<0.001. (C) Upper: immunofluorescence images of aberrant mitosis from MCF-7 cells with control (Con), YAP overexpression (YAP) or YAP overexpression plus TEAD4 knocking down (YAP+siTEAD4). Lower: immunofluorescence images of aberrant mitosis from MDA-MB-231 transfected with scramble siRNA (siNC) or siTEAD4 siRNA (siTEAD4-1#, TEAD4-2#). β-Tubulin (red), γ-Tubulin (green), Nuclei (blue). Scale bar: 10μm. Histograms showed the mean percentage ± SD of aberrant mitosis rate. ns, no significant; **p<0.01; ***p<0.001. (D) Upper: images of chromosome metaphase spreading in MCF-7 cells with control (Con), YAP overexpression (YAP) or YAP overexpression plus TEAD4 knocking down (YAP+siTEAD4). Lower: images of chromosome metaphase spreading in MDA-MB-231 transfected with scramble siRNA (siNC) or siTEAD4 siRNA (siTEAD4-1#, TEAD4-2#). Scale bar: 5μm. Histograms showed the mean percentage ± SD of cell rate with aberrant chromosomal number. ns, no significant; **p<0.01; ***p<0.001. (E) MCF-7 control (Con) or YAP overexpression (YAP) cells were treated with DMSO, Verteporfin (at a dose of 1μM) or TED-347 (at a dose of 10μM) for 16 hours, respectively. Aberrant mitosis was observed by immunofluorescence. β-Tubulin (red), γ-Tubulin (green), Nuclei (blue), scale bar: 10μm. Histograms show the mean percentage ± SD of aberrant mitosis rate. ns, no significant; **p<0.01. (F) MDA-MB-231 cells were treated with DMSO, Verteporfin (at a dose of 1μM) or TED-347 (at a dose of 10μM) for 16 hours, respectively. Aberrant mitosis was observed by immunofluorescence. β-Tubulin (red), γ-Tubulin (green), Nuclei (blue), scale bar: 10μm. Histograms show the mean percentage ± SD of aberrant mitosis rate. ns, no significant; **p<0.01. (G) MCF-7 control (Con) or YAP overexpression (YAP) cells were treated with DMSO, Verteporfin (at a dose of 1μM) or TED-347 (at a dose of 10μM) for 24 hours, respectively. Karyotype analysis was performed via chromosome metaphase spreading. scale bar: 5μm. Histograms showed the mean percentage ± SD of cell rate with aberrant chromosomal number. ns, no significant; *p<0.05; and ***p<0.001. (H) MDA-MB-231 cells were treated with DMSO, Verteporfin (at a dose of 1μM) or TED-347 (at a dose of 10μM) for 24 hours, respectively. Karyotype analysis was performed via chromosome metaphase spreading. scale bar: 5μm. Histograms showed the mean percentage ± SD of cell rate with aberrant chromosomal number. *p<0.05; and **p<0.01.

### YAP/TEAD transcriptionally activating spindle assembly checkpoint AJUBA expression

To identify the downstream targets of YAP-TEADs mediated CIN, we re-analyzed the published ChIP sequence data of YAP/TEAD and YAP/TAZ expression profile in MDA-MB-231 from GSE66081 dataset^18^. Combining with ChIP sequencing and expression profile, we found 3987 genes were co-interacted with both YAP and TEAD4, among them, 205 genes were differential expression in cells with YAP/TAZ knocking down (Fig. 3A). Thus, these genes (102 were up-regulated and 103 were down-regulated via siYAP/TAZ) appeared to be potential downstream target and were selected for further study (Fig. 3B). Gene ontology analysis revealed that the down-regulated genes could enriched into “mitotic cycle process” category (15 genes were included), which involved in aberrant mitosis (Fig. 3C). To further determine the exact downstream, RT-qPCR was performed to conform the expression level of all these 15 genes in MDA-MB-231 with YAP knocking down, and the results showed that AJUBA exhibited the most significant expression fold change (Fig. S3A).

**Figure 3.**
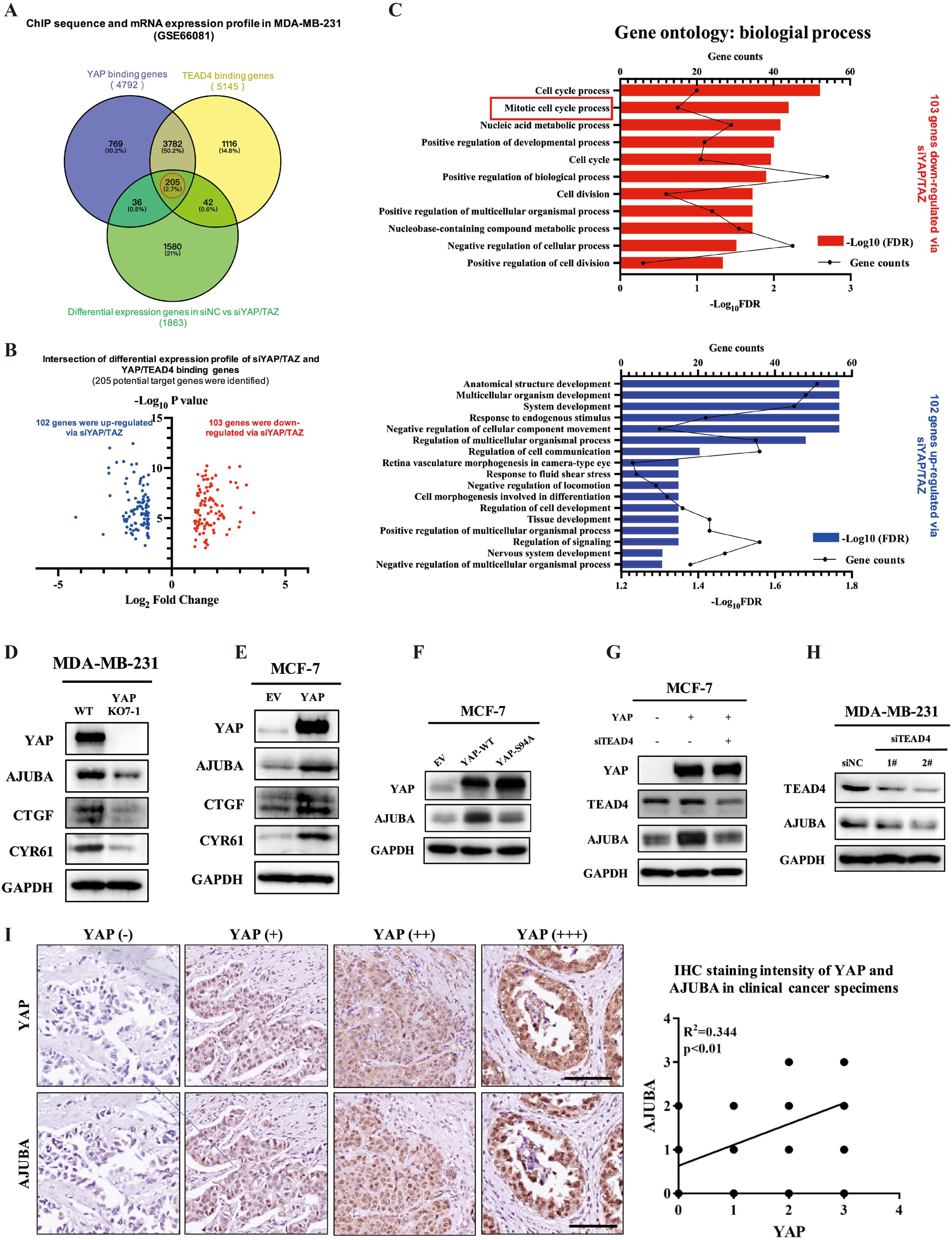
YAP/TEAD transcriptionally activating spindle assembly checkpoint AJUBA expression. (A) Venn diagram indicated the intersection between YAP/TEAD binding genes and the differential expression genes of siNC vs. siYAP/TAZ in MDA-MB-231 cells. Data were obtained from GSE66081. 205 intersected genes (red circle) were collected for further analysis. (B) Volcano diagram showed the intersected genes from (A). 102 genes (blue dots) were upregulated, while 103 genes (red dots) were downregulated via siYAP/TAZ. (C) Gene ontology enrichment of the 102 up-regulated genes (upper) and 103 down-regulated genes (lower) indicated in (B). Biological processes involved in molecular function were presented. Bar plots represent -Log_10_ false discovery rate (FDR), and line charts showed gene counts. (D) Cell lysates from MDA-MB-231 with wild type (WT) and YAP knocking out (YAP KO7-1) cell lines were used for immunoblotting. Lysates were probed for YAP, AJUBA, CTGF, CYR61, and GAPDH was used as a loading control. (E) Cell lysates from MCF-7 cells transfected with empty vector (EV) or YAP overexpressing plasmids (YAP) were used for immunoblot. Lysates were probed for YAP, AJUBA, CTGF, CYR61, and GAPDH was used as a loading control. (F) Cell lysates from MCF-7 cells transfected with empty vector (EV), wild type YAP (YAP-WT), or YAP S94A mutant (YAP-S94A) were used for immunoblot. Lysates were probed for YAP, AJUBA, and GAPDH was used as a loading control. (G) Cell lysates from MCF-7 cells transfected with YAP overexpressing plasmid (YAP) and/or siTEAD4 siRNA (siTEAD4) were used for immunoblot. Empty vector and scramble siRNA (siNC) was used as negative control. Lysates were probed for YAP, AJUBA and TEAD4, and GAPDH was used as a loading control. (H) Cell lysates from MDA-MB-231 cells transfected with siTEAD4 (siTEAD4-1# and siTEAD4-2#) were collected for immunoblot. Scramble siRNA (siNC) was used as negative control. Lysates were probed for TEAD4, AJUBA, and GAPDH was used as a loading control. (I) Representative images of YAP and AJUBA IHC staining from tumors with different YAP expression level in human pan-cancer specimen microarray (scale bar: 250μm). YAP (−): YAP IHC intensity= 0, n=26; YAP (+): YAP IHC intensity =1, n=95; YAP (++): YAP IHC intensity=2, n=47; and YAP (+++): YAP IHC intensity=3, n=22. The correlation between YAP and AJUBA IHC staining intensity was evaluated via IHC and calculated by Pearson analysis, R^2^=0.344 and p<0.01.

AJUBA, termed as a LIM protein, could act as a versatile scaffold involving in centrosome assemble, thus regulate spindle formation and mitosis^25,26^. To confirm whether YAP could induce AJUBA expression, we first knocked out YAP in MDA-MB-231 cells via CRISPR/Cas9 and overexpressed YAP in MCF7 cells. The results showed that, knocking out YAP in MDA-MB-231 cells could significantly reduce the mRNA and protein level of AJUBA, in contrary, overexpressing YAP could significantly induce AJUBA expression in MCF7 cells (Fig. S3B, C and Fig. 3D, E). Similarly, TAZ could also positive regulate AJUBA expression in MDA-MB-231 and MCF7 (Fig. S3D, E).

Next, we validated whether TEAD was essential in AJUBA expression. In MCF7 cells, the expression of YAP-S94A mutant was unable to activate AJUBA transcription (Fig. 3F), meanwhile, either knocking down TEAD4 or inhibiting YAP-TEADs interaction could abolished YAP/TAZ induced AJUBA expression in MCF7 cells (Fig. 3G and Fig. S3F-H). Similarly, in MDA-MB-231 cells, either knocking down TEAD4 or treating with verteporfin/TED-347 could inhibit AJUBA expression in MDA-MB-231 cells (Fig. 3H, M and Fig. S3I, J). IHC staining of tissue array also showed a positive correlation between YAP and AJUBA expression in clinical tumor specimens of multiple cancer (Fig. 3I). These findings indicate that YAP/TEADs could transcriptionally activating AJUBA expression

### YAP regulates CIN through AJUBA-AURKA axis

AJUBA has been reported to be an upstream of AURKA kinase and could activate AURKA through promoting its phosphorylation at T288, then phosphorylated AURKA maintains a proper segregation of centrosomes and activates spindle assembly^27^. Western blot assay showed that YAP could induce AJUBA expression and phosphorylated AURKA level in both MDA-MB-231 and MCF7 cells (Fig.4A, B). According to previous researches, an ectopic abundance of p-AURKA could lead to multipolar mitosis and cause CIN formation^28^ To unveil whether YAP-induced CIN was caused by aberrant AJUBA-AURKA signaling, we first knocked down AJUBA in MDA-MB-231 cells and overexpressed exogenous AJUBA in MCF7 cells. Western bot assay verified that AJUBA was positively correlated with AURKA T288 phosphorylation level (Fig. S4A, B). Knocking down AJUBA could significantly reduce the proportion of cells with multipolar mitosis and aneuploidy in MDA-MB-231 cell (Fig. S4C, D), similarly, overexpressing AJUBA could significantly induce aberrant mitosis and aneuploidy in MCF7 cells (Fig. S4E, F). Next, we up-regulated AJUBA expression in MDA-MB-231 cells transfected with YAP interfering siRNAs, and found the YAP-inhibited AURKA phosphorylation could be significantly reversed by AJUBA (Fig. 4C). Knocking down AJUBA could inhibited YAP induced AURKA phosphorylation in MCF7 (Fig. 4D) and MCF10A cells (Fig. S4G). Immunofluorescence and karyotype analysis also revealed that overexpressing AJUBA could rescue aberrant mitosis cell proportion and aneuploidy rate in MDA-MB-231 cells with YAP knockdown (Fig. 4E, F), meanwhile, depletion of AJUBA by siRNAs could significantly diminished aberrant mitosis and aneuploidy in MCF7 and MCF-10A cells with YAP overexpression (Fig. 4G, H and Fig. S4H, I). Similarly, directly inhibiting AURKA phosphorylation using a high selective inhibitor Aliertib (MLN8237) could also reverse YAP induced AURKA T288 phosphorylation, multipolar mitosis and aneuploidy in both MCF7 (Fig. 4I-K) and MCF10A (Fig. S4J-L). These data indicate that YAP could regulate CIN via AJUBA-AURKA axis.

**Figure 4.**
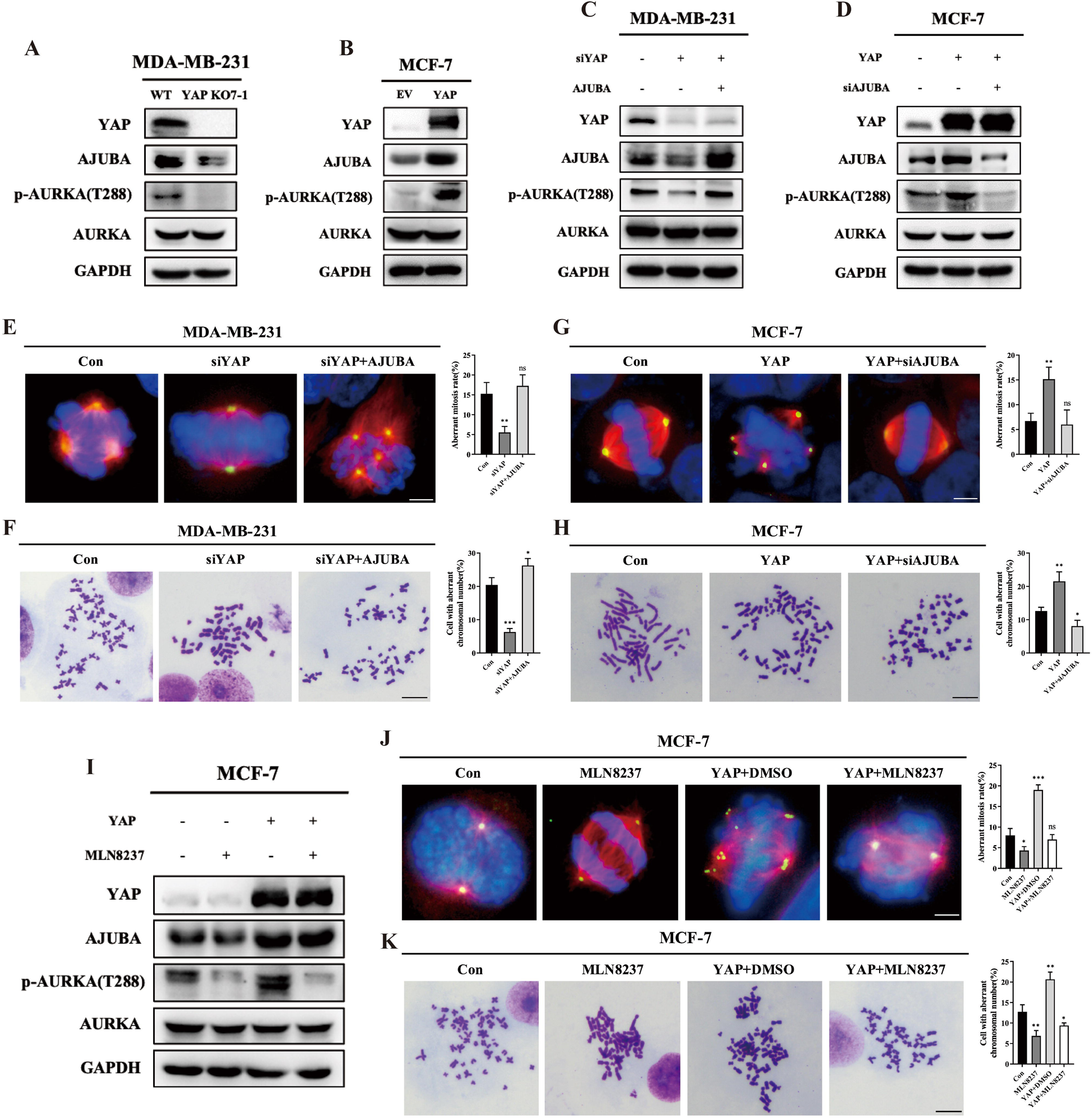
YAP regulates CIN through AJUBA-AURKA axis. (A) Cell lysates from MDA-MB-231 with wild type (WT) and YAP knocking out (YAP KO7-1) cell lines were used for immunoblot. Lysates were probed for YAP, AJUBA, AURKA, p-AURKA (T288). GAPDH was used as a loading control. (B) Cell lysates from MCF-7 cells transfected with empty vector (EV) or YAP overexpressing plasmid (YAP) were used for immunoblot. Lysates were probed for YAP, AJUBA, AURKA, p-AURKA (T288). GAPDH was used as a loading control. (C) Cell lysates from MDA-MB-231 cells transfected with siYAP siRNAs (siYAP) and/or AJUBA overexpressing plasmids (AJUBA) were collected for immunoblot. Empty vector and scramble siRNA (siNC) was used as negative control. Lysates were probed for YAP, AJUBA, AURKA, p-AURKA (T288). GAPDH was used as a loading control. (D) Cell lysates from MCF-7 cells transfected with YAP overexpressing plasmid (YAP) and/or siAJUBA siRNAs (siAJUBA) were collected for immunoblot. Empty vector and scramble siRNA (siNC) was used as negative control. Lysates were probed for YAP, AJUBA, AURKA, p-AURKA (T288), and GAPDH was used as a loading control. (E) Representative images of aberrant mitosis from MDA-MB-231 cells transfected with siYAP siRNAs and/or AJUBA overexpressing plasmids (AJUBA). Empty vector and scramble siRNA (siNC) was used as negative control. β-Tubulin (red), γ-Tubulin (green), Nuclei (blue). Scale bar: 10μm. Histograms showed the mean percentage ± SD of aberrant mitosis rate. ns, no significant difference; **p<0.01. (F) Representative images of chromosome metaphase spreading from MDA-MB-231 cells transfected with siYAP siRNAs and/or AJUBA overexpressing plasmids (AJUBA). Empty vector and scramble siRNA (siNC) was used as negative control. Scale bar: 5 μm. Histograms show the mean percentage ± SD of cell rate with aberrant chromosomal number. *p<0.05; ***p<0.001. (G) Representative images of aberrant mitosis from MCF-7 cells transfected with YAP overexpressing plasmid (YAP) and/or siAJUBA siRNAs (siAJUBA). Empty vector and scramble siRNA (siNC) was used as negative control. β-Tubulin (red), γ-Tubulin (green), Nuclei (blue). Scale bar: 10μm. Histograms showed the mean percentage ± SD of aberrant mitosis rate. ns, no significant difference. ns, no significant difference; **p<0.01. (H) Representative images of chromosome metaphase spreading from MCF-7 cells transfected with YAP overexpressing plasmid (YAP) and/or siAJUBA siRNAs (siAJUBA). Empty vector and scramble siRNA (siNC) was used as negative control. Scale bar: 5 μm. Histograms show the mean percentage ± SD of cell rate with aberrant chromosomal number. *p<0.05; **p<0.01. (I) MCF-7 cells transfected with empty vector or YAP overexpressing plasmids (YAP) were treated with DMSO or MLN8237 at 10 μM for 24 h. Protein level of YAP, AJUBA, AURKA and p-AURKA (T288) were examined via western blot. GAPDH was used as a loading control. (J) MCF-7 cells with control (Con) or YAP overexpression (YAP) were treated with DMSO or MLN8237 at a dose of 10μM for 16 hours. Aberrant mitosis was observed by immunofluorescence. β-Tubulin (red), γ-Tubulin (green), Nuclei (blue), scale bar: 10μm. Histograms show the mean percentage ± SD of aberrant mitosis rate. ns, no significant. ns, no significant difference; *p<0.05; ***p<0.001. (K) MCF-7 cells with control (Con) or YAP overexpression (YAP) were treated with DMSO or MLN8237 at a dose of 10μM for 16 hours. Karyotype analysis was performed via chromosome metaphase spreading. scale bar: 5μm. Histograms showed the mean percentage ± SD of cell rate with aberrant chromosomal number. ns, no significant; *p<0.05; **p<0.01 and ***p<0.001.

### YAP/TEAD promotes AJUBA transcription through its super enhancer

To elucidate the detailed mechanism of YAP/TEADs induced AJUBA transcription, we annotated the published ChIP-sequence data of MCF7 cells in ENCODE database, and identified three TEAD4 binding peaks on AJUBA genome. Among them, one peak located in the conventional promoter region (−2000bp to +50bp TSS), and the remaining two peaks (named as E1 and E2) was overlapped with the H3K27ac binding region, which was defined as potential super enhancer region by ROSE software (Fig. 5A). Next, we validated that YAP and TEAD4 could also combined with the promoter and E1, E2 region of AJUBA gene in MDA-MB-231 cells using chromatin immunoprecipitation (Fig. 5B). To evaluate the transcriptional activity of YAP-TEAD4 on AJUBA gene, we transfected YAP and its mutants (S127A: continuous active mutant; S94A: TEAD domain defected mutant) into HEK293T cell (Fig. S5A) and performed Dual-Luciferase reporter assay containing E1/E2 and AJUBA promoter sequence. The results showed that the transcription of E1/E2 and AJUBA promoter could be significantly activated by wild type YAP and S127A mutant, while luciferase activity of all these reporters were relatively limited in YAP-S94A group (Fig. 5C). Moreover, mutated TEAD4 motif in E1/E2 and promoter sequences could abolish YAP induced transcriptional activity (Fig. 5D). To further investigate the function of E1/E2 in AJUBA transcription, we synthesized two sgRNAs targeting the E1 (sgRNA-1) and E2 (sgRNA-2) and verified their efficiency in MCF7 (Fig. S5B, C). Knocking out E1/E2 via CRISPR-CAS9 technique could also abolish YAP induced AJUBA transcription in MCF7 cells (Fig. 5E). CRISPR activation/interference assay also showed that activating E1/E2 could significantly induce AJUBA transcription in MCF7 (Fig. 5F), while interfering E1/E2 could in reverse decrease AJUBA transcription level in MDA-MB-231 (Fig. 5G). Interesting, we found E1/E2 could amplify the transcription activity of AJUBA promoter in HEK-293T cells with YAP overexpression (Fig. 5H). Therefore, we indicated that YAP/TEAD could interact with AJUBA super enhancer region, thus activated its promoter and induce its transcription in breast cancer.

**Figure 5.**
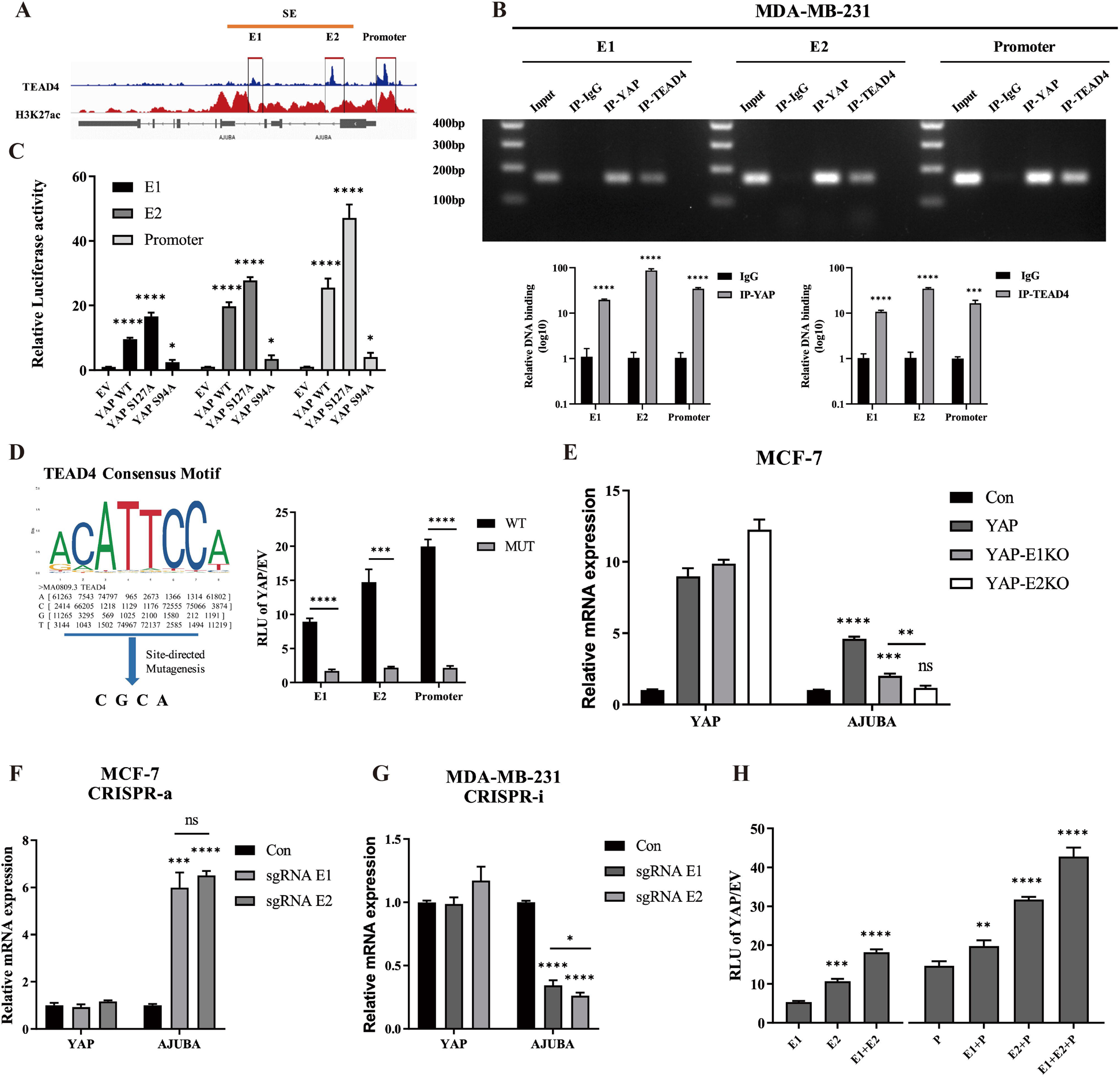
YAP/TEAD promotes AJUBA transcription through its super enhancer. (A) ChIP-seq track diagram of H3K27ac and TEAD4 on AJUBA locus. Data were obtained from ENCODE database. SE: Super enhancer; E1: Enhancer 1; E2: Enhancer 2. (B) ChIP assay was performed using YAP or TEAD4 antibodies in MDA-MB-231 cells. IgG antibody was used as negative control. DNA in the immunocomplex was amplified with the primers targeting E1, E2 or AJUBA promoter, and measured via agarose gel electrophoresis (upper) or qPCR (lower). ***p<0.001 and ****p<0.0001. (C) Dual Luciferase reporter assay was performed to evaluate the relative transcriptional activity of E1, E2 or AJUBA promoter in HEK-293T cells transfected with empty vector (EV), wild type YAP (YAP WT), YAP S127A mutant (YAP S127A), or YAP S94A mutant (YAP S94A). *p<0.05 and ****p<0.0001. (D) Left: diagram of site-direct mutagenesis on TEAD4 motif. TEAD4 motif sequence was obtained from JASPAR database. Right: Dual Luciferase report assay was performed on HEK-293T cells with empty vector (EV) or YAP overexpression. The transcriptional activity of E1, E2 and AJUBA promoter with wild type or mutated TEAD4 motif was evaluated by ratio of RLU in cells with YAP overexpression to RLU in cells with EV overexpression. RLU, relative light unit. ***p<0.001 and ****p<0.0001. (E) AJUBA mRNA expression level in MCF-7 with control (MCF-7 Con), YAP overexpression (MCF-7 YAP), YAP overexpression plus E1 knockout (YAP-E1KO) and YAP overexpression plus E2 knockout (YAP-E2KO). GAPDH was used as an endogenous control. ns. no significant difference; **p<0.01; ***p<0.001 and ****p<0.0001. (F) CRISPR activation assay (CRISPR-a) was performed in MCF7 cells using sgRNA control (Con), sgRNAs targeting E1 (sgRNA E1) or sgRNAs targeting E2 (sgRNA E2) sequence on AJUBA locus. The AJUBA mRNA expression level was quantified via qPCR assay. GAPDH was used as an endogenous control. ns. no significant difference; ***p<0.001 and ****p<0.0001. (G) CRISPR interference assay (CRISPR-i) was performed in MDA-MB-231 cells using sgRNA control (Con), sgRNAs targeting E1 (sgRNA E1) or sgRNAs targeting E2 (sgRNA E2) sequence on AJUBA locus. The AJUBA mRNA expression level was quantified via qPCR assay. GAPDH was used as an endogenous control. *p<0.05; ***p<0.001 and ****p<0.0001. (H) Dual Luciferase reporter assay was performed on HEK-293T cells with empty vector (EV) or YAP overexpression. The transcriptional activity of E1, E2, AJUBA promoter (P), E1+E2, E1+P, E2+P and E1+E2+P with wild type or mutated TEAD4 motif was evaluated. The transcriptional activity was normalized by the ratio of RLU in cells with YAP overexpression to RLU in cells with EV overexpression. RLU, relative light unit. **p<0.01; ***p<0.001 and ****p<0.0001.

### Super enhancer is vital for YAP induce AJUBA expression and CIN formation

To further validate the crucial role of super enhancer in YAP induced AJUBA transcription and CIN formation, we first used JQ1, an unselective BET-inhibitor^29^, to inhibit widespread enhancer activity in MCF7 and MDA-MB-231 cells. The results showed that an increasement of JQ-1 concentration, the protein level of AJUBA was gradually declined (Fig. 6A, B). Using CRISPR activation/interference assay, we found activating E1/E2 could significantly induce AJUBA expression in MCF7 (Fig. 6C). In contrast, interfering E1/E2 could decrease AJUBA protein level in MDA-MB-231 cells (Fig. 6D). Immunofluorescence and karyotype analysis also showed that activating E1/E2 could in reverse induce aberrant mitosis and aneuploidy formation in MCF7 (Fig. 6E, F), while interfering E1/E2 could inhibit multipolar mitosis and decrease aneuploidy in MDA-MB-231 (Fig. 6G, H). Furthermore, we deleted E1 and E2 in MCF7 cells via CRISPR/Cas9 technique. In cells with E1/E2 deletion, the increasement of AJUBA protein level via YAP overexpressing was partially abolished (Fig. 6I). Moreover, deleting E1/E2 could also reverse YAP induced aberrant mitosis and aneuploidy formation (Fig. 6J, K).

**Figure 6.**
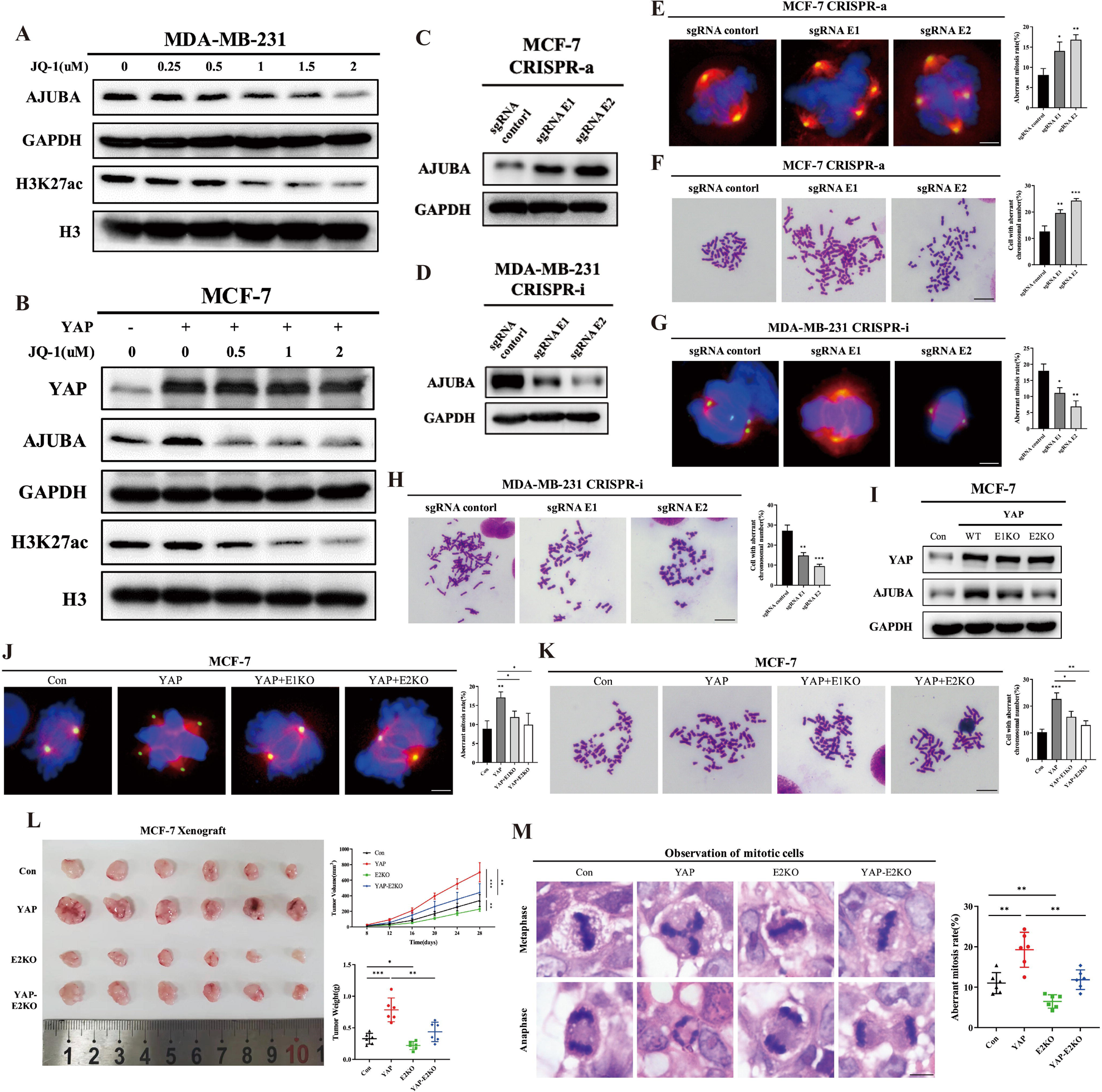
Super enhancer is vital for YAP induce AJUBA expression and CIN formation. (A) MDA-MB-231 cells were treated with indicated concentration of JQ-1 for 24 h, then protein level of AJUBA and H3K27ac were examined via western blot. GAPDH and histone H3 (H3) were used as loading control. (B) MCF-7 cells with control or YAP overexpression were treated with indicated concentration of JQ-1 for 24 h. DMSO was used as negative control. The protein level of YAP, AJUBA and H3K27ac were examined via western blot. GAPDH and H3 were used as loading control. (C) CRISPR activation assay (CRISPR-a) was performed in MCF7 cells using sgRNA control, sgRNAs targeting E1 (sgRNA E1) or sgRNAs targeting E2 (sgRNA E2) on AJUBA locus. The protein level of AJUBA were examined via western blot. GAPDH was used as loading control. (D) CRISPR interference assay (CRISPR-i) was performed in MDA-MB-231 cells using sgRNA control, sgRNAs targeting E1 (sgRNA E1) or sgRNAs targeting E2 (sgRNA E2) on AJUBA locus. The protein level of AJUBA were examined via western blot. GAPDH was used as loading control. (E) CRISPR-activation was performed in MCF7 cells using sgRNA control, sgRNAs targeting E1 (sgRNA E1) or sgRNAs targeting E2 (sgRNA E2) on AJUBA locus. Aberrant mitosis was observed by immunofluorescence. β-Tubulin (red), γ-Tubulin (green), Nuclei (blue), scale bar: 10μm. Histograms show the mean percentage ± SD of aberrant mitosis rate. ns, no significant. *p<0.05; **p<0.01. (F) Representative images of chromosome metaphase spreading from MCF-7 cells treated with CRISPR-activation of sgRNA control, sgRNAs targeting E1 (sgRNA E1) or sgRNAs targeting E2 (sgRNA E2). Scale bar: 5 μm. Histograms show the mean percentage ± SD of cell rate with aberrant chromosomal number. **p<0.01; ***p<0.001. (G) CRISPR-interference was performed in MDA-MB-231 cells using sgRNA control, sgRNAs targeting E1 (sgRNA E1) or sgRNAs targeting E2 (sgRNA E2) on AJUBA locus. Aberrant mitosis was observed by immunofluorescence. β-Tubulin (red), γ-Tubulin (green), Nuclei (blue), scale bar: 10μm. Histograms show the mean percentage ± SD of aberrant mitosis rate. ns, no significant. *p<0.05; **p<0.01. (H) Representative images of chromosome metaphase spreading from MDA-MB-231 cells treated with CRISPR-interference of sgRNA control, sgRNAs targeting E1 (sgRNA E1) or sgRNAs targeting E2 (sgRNA E2). Scale bar: 5 μm. Histograms show the mean percentage ± SD of cell rate with aberrant chromosomal number. **p<0.01; ***p<0.001. (I) AJUBA protein level in MCF-7 with control (MCF-7 Con), wild type YAP overexpression (MCF-7 YAP-WT), wild type YAP overexpression plus E1 knockout (YAP-E1KO) and YAP overexpression plus E2 knockout (YAP-E2KO) were examined via western blot. GAPDH was used as an endogenous control. (J) Representative images of Aberrant mitosis in MCF-7 cells with control (MCF-7 Con), YAP overexpression (MCF-7 YAP), YAP overexpression plus E1 knockout (YAP-E1KO) and YAP overexpression plus E2 knockout (YAP-E2KO). β-Tubulin (red), γ-Tubulin (green), Nuclei (blue), scale bar: 10μm. Histograms show the mean percentage ± SD of aberrant mitosis rate. ns, no significant. *p<0.05; **p<0.01. (K) Representative images of chromosome metaphase spreading from MCF-7 cells with control (MCF-7 Con), YAP overexpression (MCF-7 YAP), YAP overexpression plus E1 knockout (YAP-E1KO) and YAP overexpression plus E2 knockout (YAP-E2KO). Scale bar: 5 μm. Histograms show the mean percentage ± SD of cell rate with aberrant chromosomal number. *p<0.05; **p<0.01; ***p<0.001. (L) MCF-7 stable cell lines with control (MCF-7 Con), YAP overexpression (MCF-7 YAP), E2 knockout (E2KO) and YAP overexpression plus E2 knockout (YAP-E2KO) were orthotopically implanted into BALB/c nude mice (n=6). Tumor growth curve and tumor weights were measured as indicated. Data were presented as mean ± SD. *p<0.05; **p<0.01; ***p<0.001. (M) H&E staining of aberrant mitosis in the xenografts from (L). Scale bar: 4μm. Scatterplot shows the mean percentage ± SD of aberrant mitosis rate. **p<0.01.

Due to the relatively high efficacy, we further test the biological function of E2 in YAP overexpression tumor in vivo. Xenograft models of MCF cells with control, YAP overexpression, E2 knockout, and YAP overexpression combined with E2 knockout were established on female NOD/SCID mice. Four weeks after tumor implantation, tumors were collected and analyzed. Tumor size measurement showed that knocking out E2 could significantly inhibit tumor growth in MCF7 cells with YAP overexpression (Fig. 6L). The proportion of aberrant mitosis was also remarkably decreased xenografted tumor with E2 knockout (Fig.6M). Taken together, we conjectured that AJUBA super enhancer region, E1/E2, was essential in YAP induced AJUBA transcription and CIN formation.

### YAP/TEAD directly activates super enhancer and induces SE-promoter proximity on AJUBA gene

To further explore how super enhancer involved in YAP induce AJUBA expression, we first used CRISPR affinity purification in situ of regulatory elements (CAPTURE) technique to uncover the interaction between YAP/TEAD and the super enhancer region of AJUBA. CAPTURE was a new technique to unbiasedly identify locus-specific chromatin-regulating protein complexes and long-range DNA interactions^30,31^. In this current research, we conjugated eGFP to biotinylated deactivated CAS9 (dCAS9) and designed sgRNA target E1/E2 region (sgRNA E1/E2) in AJUBA super enhancer (Fig. S6A). Immunofluorescence showed that sgRNAs targeting E1 and E2 were able to recruit eGFP fused dCAS9 in MCF7 cells (Fig. S6B). Immunoprecipitation and western blot analysis showed that sgRNA E1/E2 linked dCas9 could pull down YAP/TEAD4, together with RNA polymerase II (POLR2A) and enhancer markers (H3K27ac, EP300, BRD4) in MDA-MB-231 cells (Fig.7A). Additionally, overexpressing wild type YAP, rather than S94A mutant, could remarkably increase the enrichment of TEAD4, as well as EP300, BRD4, POLR2A and acetylated H3K27 in super enhancer region of AJUBA (Fig.7B). Promoter and enhancer could form spatial loop to initiate transcription^32^, thus we determined whether YAP could enhance the interaction between super enhancer and promoter on AJUBA genome. Hi-C analysis of ENCODE dataset and CAPTURE assay revealed potential chromatin-chromatin interactions between promoter and E1/E2 on AJUBA gene locus (Fig.7C, D). Meanwhile, overexpressing wild type YAP could significantly induce the interaction between E1/E2 and promoter in MCF7 cells (Fig.7E).

**Figure 7.**
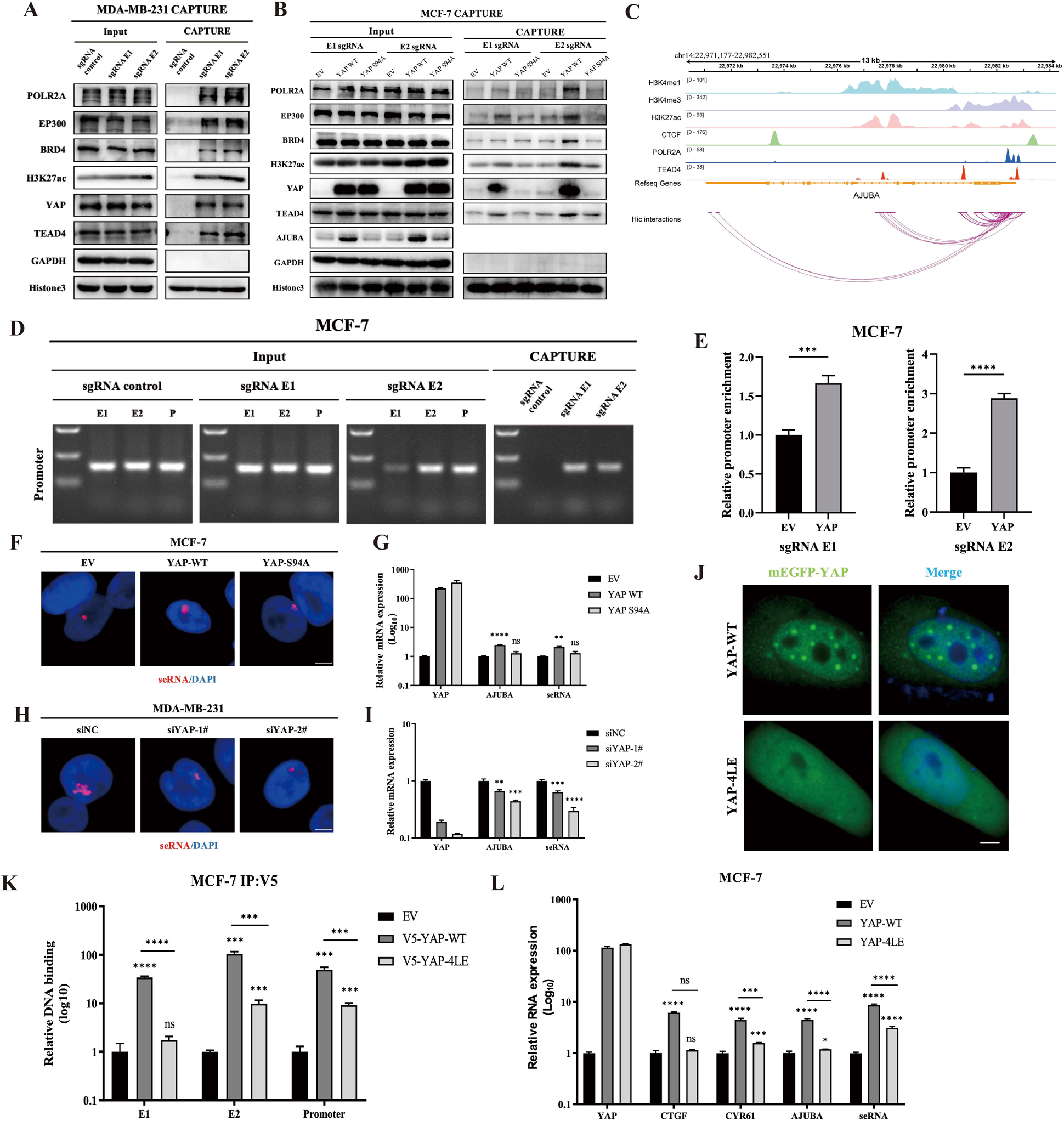
YAP/TEAD directly activates super enhancer and induces SE-promoter proximity on AJUBA gene. (A) CAPTURE assay was performed to evaluate the potential binding proteins on E1/E2 locus of AJUBA gene in MDA-MB-231 cells. Polycistronic sgRNAs targeting E1/E2 were used in the immunoprecipitation; and the protein level of POLR2A, EP300, BRD4, H3K27ac, YAP, and TEAD4 were examined via western blot. GAPDH and Histone were used as loading control. (B) CAPTURE assay was performed to evaluate the potential binding proteins on E1/E2 locus of AJUBA gene in MCF7 cells with empty vector (EV), wild type YAP (YAP WT) or YAP-S94A mutant (YAP S94A) overexpression. Polycistronic sgRNAs targeting E1/E2 were used in the immunoprecipitation; and the protein level of POLR2A, EP300, BRD4, H3K27ac, YAP, and TEAD4 were examined via western blot. GAPDH and Histone were used as loading control. (C) The chromatin-chromatin interactions between promoter and E1/E2 of AJUBA gene in MCF7 cells were analysis via Hi-C data from the ENCODE database, and were shown as indicated. H3K4me1, H3K4me3 and H3K27ac were used to mark super enhancer. POLR2A was used to mark transcriptional starting site and CTCF was used to mark topologically associating domain. TEAD4 binding sites on AJUBA locus in MCF7 were also marked in the diagram. (D) CAPTURE assay was performed to validate the potential interaction between promoter and E1/E2 locus of AJUBA gene in MCF7 cells. Polycistronic sgRNAs targeting E1/E2 were used in the immunoprecipitation. The input and affinity-captured DNA products were amplified using primers targeting AJUBA promoter and analyzed by 1% agarose gel electrophoresis. (E) CAPTURE assay and qPCR analyses were performed to detect the enrichment of in situ CAPTURE affinity-purified AJUBA promoter on E1/E2 locus in MCF-7 cells with YAP overexpression. Empty vector (EV) was used as negative control. ***p<0.001; ****p<0.0001. (F) RNA-FISH assay was performed to detect AJUBA seRNA expression in MCF-7 cells with empty vector (EV), wild type YAP (YAP WT) or YAP-S94A mutant (YAP S94A) overexpression, and the representative images were shown. Probes for seRNA were conjugated with Cy3 (red) and the nuclei were stained with DAPI. scale bar: 5 μm. (G) qPCR detecting the expression level of seRNA and mRNA of AJUBA gene in MCF-7 cells with empty vector (EV), wild type YAP (YAP WT) or YAP-S94A mutant (YAP S94A) overexpression. GAPDH was used as an endogenous control. ns, no significant. **p<0.01; ****p<0.0001. (H) RNA-FISH assay was performed to detect AJUBA seRNA expression in MDA-MB-231 cells transfected with scramble siRNA (siNC) or siYAP (siYAP-1, siYAP-2), and the representative images were shown. Probes for seRNA were conjugated with Cy3 (red) and the nuclei were stained with DAPI. scale bar: 5μm. (I) qPCR detecting the expression level of seRNA and mRNA of AJUBA gene in MDA-MB-231 cells transfected with scramble siRNA (siNC) or siYAP (siYAP-1, siYAP-2). GAPDH was used as an endogenous control. **p<0.01; ***p<0.001; ****p<0.0001. (J) mEGFP fused wild type YAP (YAP WT) or mEGFP fused YAP mutant (YAP 4LE) were transiently transfected into MCF-7 cells and the representative images of YAP protein phase separation were shown. Scale bar: 5μm. (K) The binding capacity of wild type YAP (V5-YAP-WT) or 4LE mutant (V5-YAP-4LE) to E1/E2 or promoter on AJUBA gene in MCF7 cells were evaluated via ChIP assay. Empty vector was used as negative control. ns., no significant difference; ***p<0.001; ****p<0.0001. (L) qPCR detecting CTCF, CYR61, AJUBA and seRNA expression in MCF7 cells with YAP wild type (YAP-WT) or 4LE mutant (4LE) overexpression. GAPDH was used as an endogenous control. ns., no significant difference; *p<0.05; ***p<0.001; ****p<0.0001.

Next, we validated whether YAP could directly activate AJUBA super enhancer. As previous reports, SE could transcribe into a non-coding RNA, termed as seRNA, when it was activated^33^. Therefore, expression level of seRNA could represent the activity of super enhancer^34^. Combining the data from FANTOM5 database (Fig. S6C) and RACE technique (Fig. S6D), we designed probes to detected AJUBA seRNA. RNA-FISH assay and RT-qPCR showed that overexpressing wild type YAP rather S94A mutant could significantly induce seRNA transcription in MCF7 (Fig.7F, G), while knocking down YAP could significantly reduce the expression level of AJUBA seRNA in MDA-MB-231 cells (Fig.7H, I).

Previous study demonstrated that YAP could form liquid-liquid phase separation (LLPS) in nucleus and enhance downstream genes expression^35^. In our research, we also observed exogeneous eGFP-YAP could form droplet-like condensates in HEK293T, which presented LLPS characteristics in 1,6-HEX assay (Fig. S6E) and FRAP assay (Fig. S6F). Interesting, when we transfected mCherry-tagged YAP into HEK-293T cells in which the super enhancer was labelled by CAPTURE (green), the YAP condensates were co-located with sgRNA E1/E2 marked region (Fig.S6G). As previous reported, coil-coil (C-C) domain was essential in YAP phase separation^35^, therefore a YAP C-C domain mutant (4LE^35^) was used in this research. The results showed that 4LE mutant could not only abolish LLPS of YAP protein (Fig.7J), but also reverse YAP induced E1/E2-promoter interaction (Fig.7K), seRNA transcription (Fig.7L) and AJUBA expression (Fig.S6H) in MCF7 cells. These data indicated that YAP could occur LLPS in the AJUBA SE region, thus enhancing SE-promoter interaction and AJUBA transcription.

## Discussion

Chromosomal instability is a major source of large-scale losses, gains and rearrangements of DNA on genome wide, therefore the broad genomic complexity caused by CIN is regarded as the hallmark of cancer^7^. Evidences show that CIN are acted as a driver of tumor heterogeneity and evolution^36^, and often positively correlated with malignant progression, chemotherapy resistance and poor prognosis^37,38^. Till now, various researchers’ effort to unveil its underlying mechanisms, however, the etiology of chromosomal instability remains uncertain.

In our current research, we observed that YAP/TAZ could directly induce aberrant mitosis in breast cancer cells, thus cause abnormal chromosomal segregation and aneuploidy. In malignancy, aneuploidy often leads to a continual generation of new chromosomal alterations, thus causes chromosomal instability and the development of intratumor heterogeneity^1,39^. Several mechanisms underlying CIN have been proposed, including weakened spindle checkpoint signaling, supernumerary centrosomes, defects in chromatid cohesion, abnormal kinetochore-microtubule attachments and increased spindle microtubule dynamics^40^. Our findings showed that overexpressing YAP/TAZ could lead to supernumerary centrosomes thus promote CIN breast cancer cells. This result revealed a novel insight of YAP/TAZ signaling in the regulation of cell mitosis in breast cancer.

Previously, Weiler SME, et al. reported that YAP could trigger aneuploid phenotype through transcriptionally activating FOXM1 expression in liver cancer^14^. However, Pan F, et al. showed that FOXM1 was in reverse upregulated upon aneuploid induction in cells and facilitate mitotic exit by inhibiting the spindle assembly checkpoint^41^. Therefore, up-regulation of FOXM1 might be the result, rather than the cause, of YAP-induced aberrant mitosis. In our study, we found YAP/TAZ could trigger CIN through inducing AJUBA expression. AJUBA has been identified as a LIM scaffold protein involving in the regulation of cell mitosis^42^. Mechanically, AJUBA could interact with AURKA, thus promoting its auto-phosphorylation at T288^43^ and activating. Phosphorated AURKA (T288) could subsequence induce the maturation and segregation of centrosomes^44^, while a hyperactivated AURKA could lead to mitotic error in tumorigenesis^45^. Interesting, some researchers reported that AJUBA could act as a upstream of Hippo pathway by inactivating Hippo pathway key kinases large tumor suppressor kinases 1/2, however, deprivation of YAP could also downregulate AJUBA expression^46,47^. Our data showed that manipulation of AJUBA had slightest influence on YAP/TAZ, at least in breast cancer cells. Therefore, a potential negative feedback circle could be existed in YAP/TAZ-AJUBA axis. Our research indicated a more direct mechanism to explain how YAP/TAZ promote chromosomal instability in breast cancer. These findings revealed a critical role of YAP/TAZ in regulation of mitosis and provide new therapeutic targets for cancer therapy.

Currently, it is not fully elucidated how YAP/TAZ pathway regulated tumor oncogenic signaling and biological process. In conventional viewpoint, the mechanism of YAP/TAZ regulating tumor progress mainly focused on interacting with transcription factors, such as TEADs, RUNX and TEFs, and activating the downstream cancer-related genes expression through its promoter, however, accumulating evidence suggested that enhancers might play much more important roles in Hippo-related tumorigenesis and progression^48^. Zhu et al. claim that YAP could act as a co-regulator on the enhancer region of estrogen-regulated genes^49^. Our previous research also revealed that YAP could induce TIAM1 expression through its enhancer and promote invadopodia formation in breast cancer^24^. Moreover, YAP/TAZ was observed to flag a large set of enhancers with super-enhancer (SE) like functional properties^21,23^. SEs could spatially interact with promoters to form the so-called SE-promoter loops to maximize the transcriptional activating effects^50^. Our results show that YAP/TEAD4 complex could form transcriptional apparatus on the super enhancer of AJUBA gene, thus strengthening the promoter-SE loop structure and boosting AJUBA transcription. Deleing or inhibiting SE of AJUBA could abolish YAP mediated aneuploid in breast cancer. Our findings demonstrated that structural and functional intact of super enhancer was essential for YAP induced AJUBA activation and CIN formation.

Intracellular liquid-liquid phase separated (LLPS) condensates have been widely investigated^51^. Considering the comparted structural nature, liquid condensate droplets are able to accommodate specific substrates and accelerate the biochemical reactions within compartment^52^. Existing evidences have revealed that transcription factors could form LLPS condensates on super enhancers region, thus strengthening the interaction between SEs and promoters^53^. In our study, we found that YAP/TEAD4 complex could occur phase separation on AJUBA-SE and the expression level of correlated seRNA was elevated. Through CAPTURE technique, BRD4 and EP300 were also observed accumulating at AJUBA-SE region, however, our present study didn’t further explore how YAP, BRD4 or other core transcription factors could directly induce phase separation in breast cancer cells. Previously, Sun et al reported the coil-coil domain of YAP which was essential in the LLPS formation^35^. The concrete domain involved in LLPS of YAP on AJUBA gene loci need to be further explored.

There are still some limitations in this study. First, we only evaluated aberrant mitosis and aneuploidy via immunofluorescence and karyotype analysis. Other key phenotypes of CIN, for example DNA-bridge and DNA lagging, were not validated in this research. Therefore, further experiment on assessing different characteristics of YAP/TAZ mediated CIN is essential in the future. Second, our results confirmed that the transcription factor, TEAD4, was necessity in YAP induced AJUBA expression and aberrant centrosome formation in breast cancer. However, whether other transcription factors, like TEF1, RUNX2, was also involved in YAP mediated mitotic error was still uncertain. Third, seRNAs were reported to regulate transcription in both cis and trans configuration, meanwhile, those located in the cytoplasm could also mediate various cell activities^34^. Although our data showed an increasement of AJUBA seRNA transcriptional level under YAP stimuli, the detailed biological function of these non-coding RNAs were still unknown. The role of seRNA in Hippo signaling should be unveil in the future.

In summary, our study unveils a novel cross-talk between Hippo and AJUBA-AURKA signaling, and provided a new insight in CIN formation in breast cancer.

## Acknowledgements

This work was supported by grants from the National Natural Science Foundation of China (Grant numbers: 82273061, 81773053) and Hubei Provincial Natural Science Foundation of China (Grant numbers: 2023AFB118).

We thank Professor Shuguo Sun (Huazhong University of Science and Technology) for providing support for phase separation examination. We thank the Experimental Medicine Center of Tongji Hospital for providing support to our experiment. We thank the Department of Pathology of Tongji Hospital for immunohistochemistry analysis of clinical specimens. We apologize to the colleagues whose work was not cited due to space constraint.

## Author contributions

J.S. and D.X. were responsible for study design and project administration. Q.H., R.Z., and J.S. were responsible for performing experiments and data collecting; R.Z., W.M., H.D., Z.Q. and Z.C. contributed to extracting and analyzing data; Q.H. and J.S. wrote this manuscript; R.Z., J.S., L.L. and D.X. revised and edited this manuscript; J.S. and D.X. contributed to funding acquisition.

## STAR Methods

### KEY RESOURCES TABLE

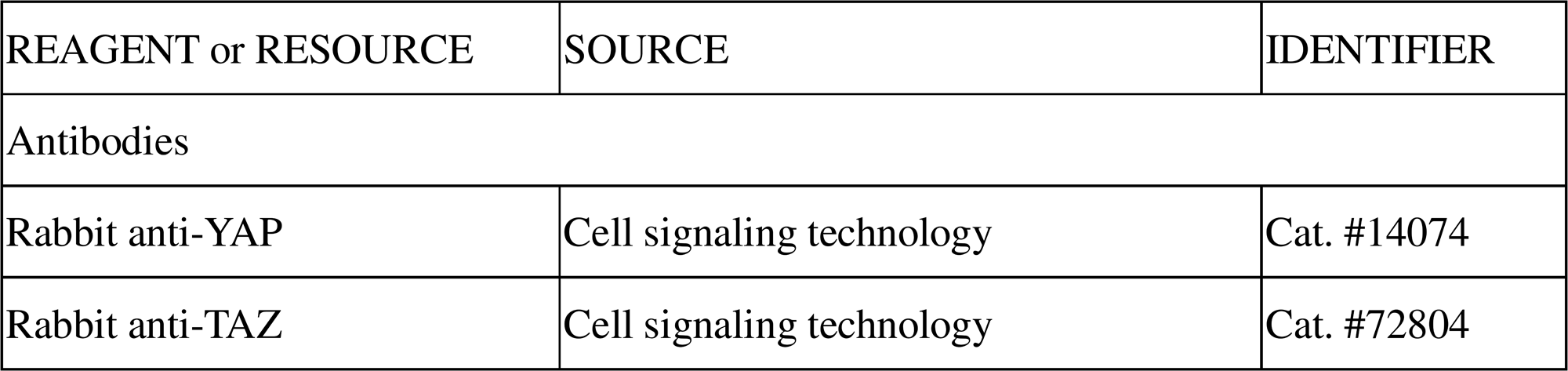
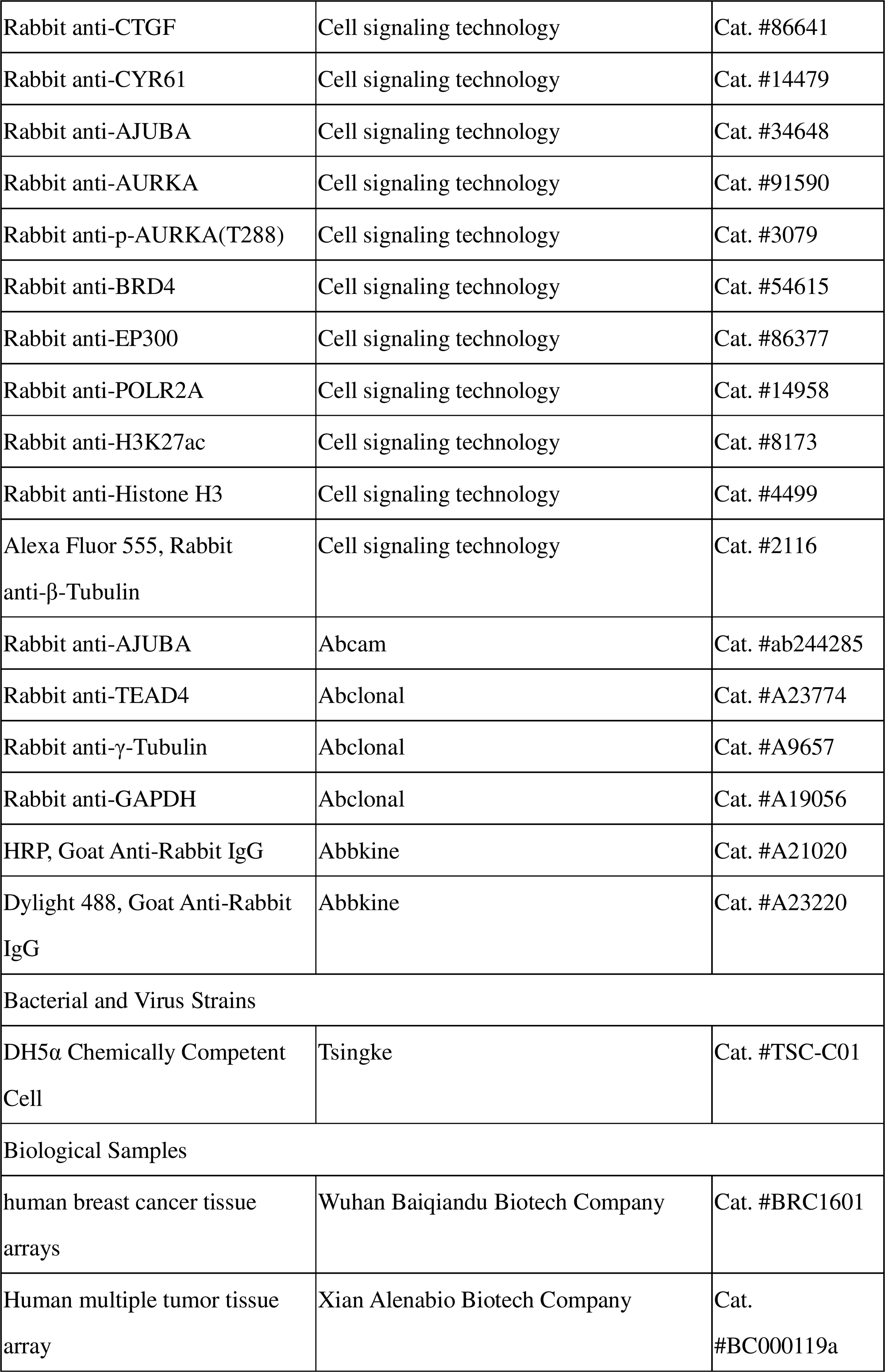
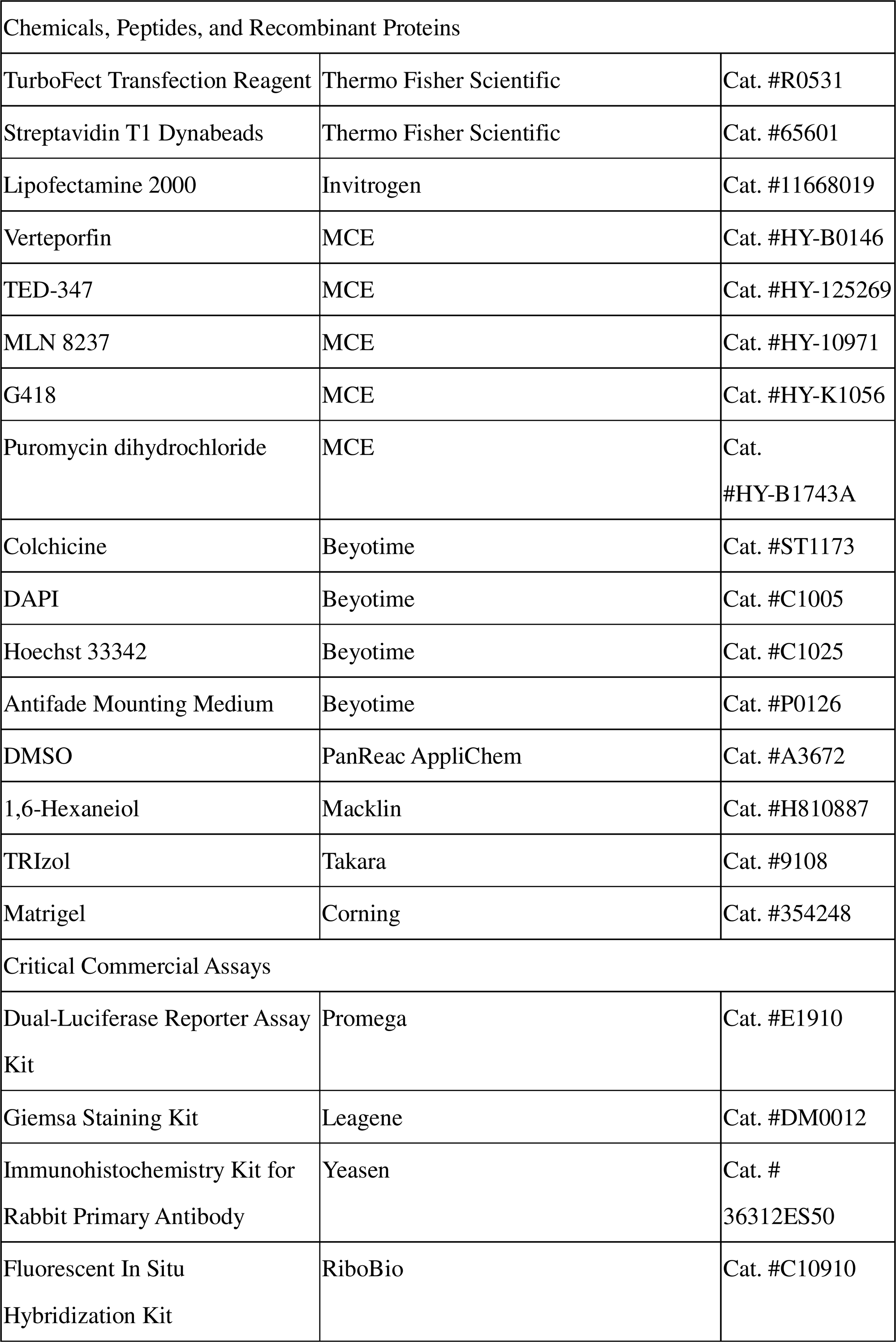
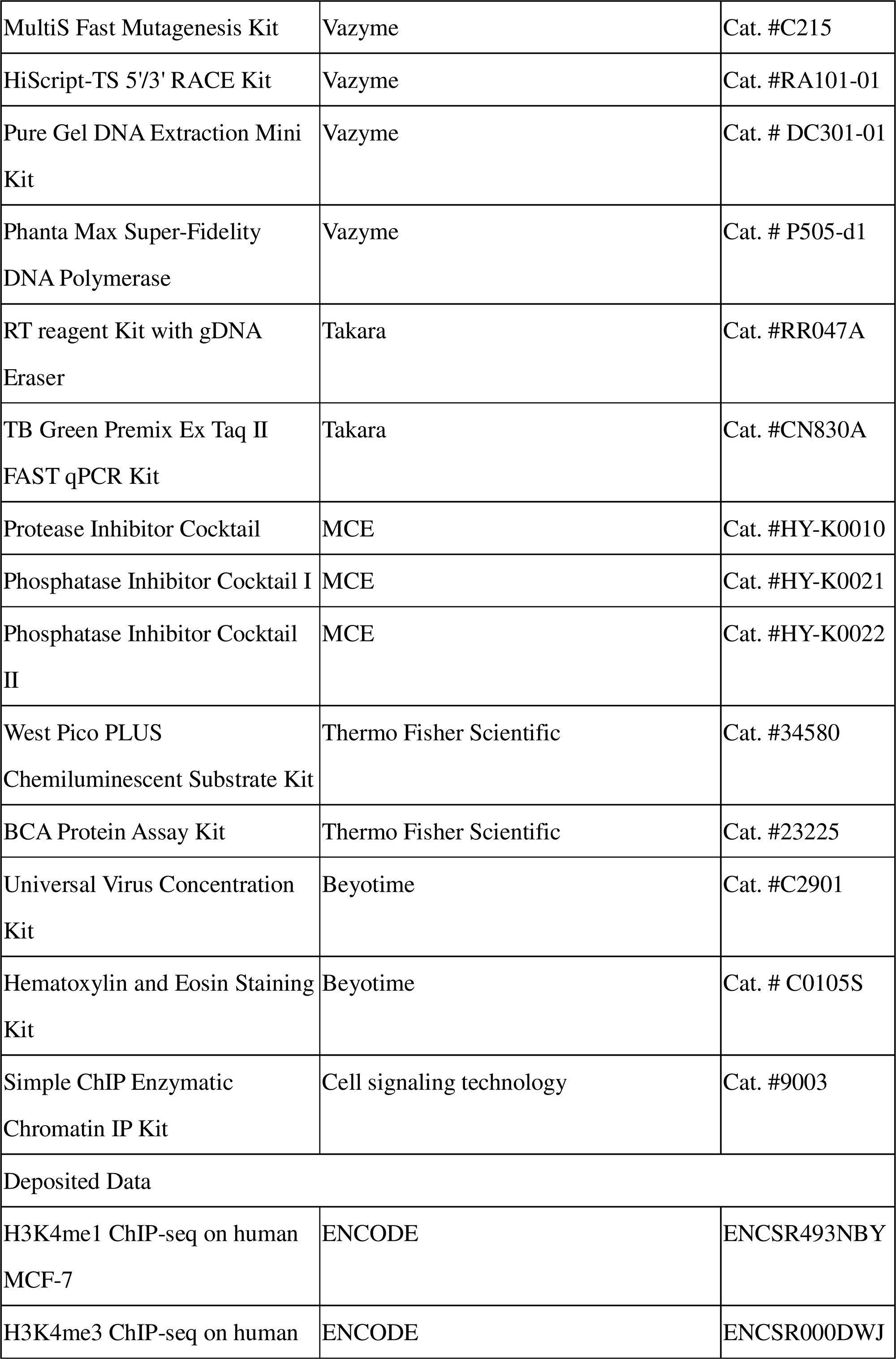
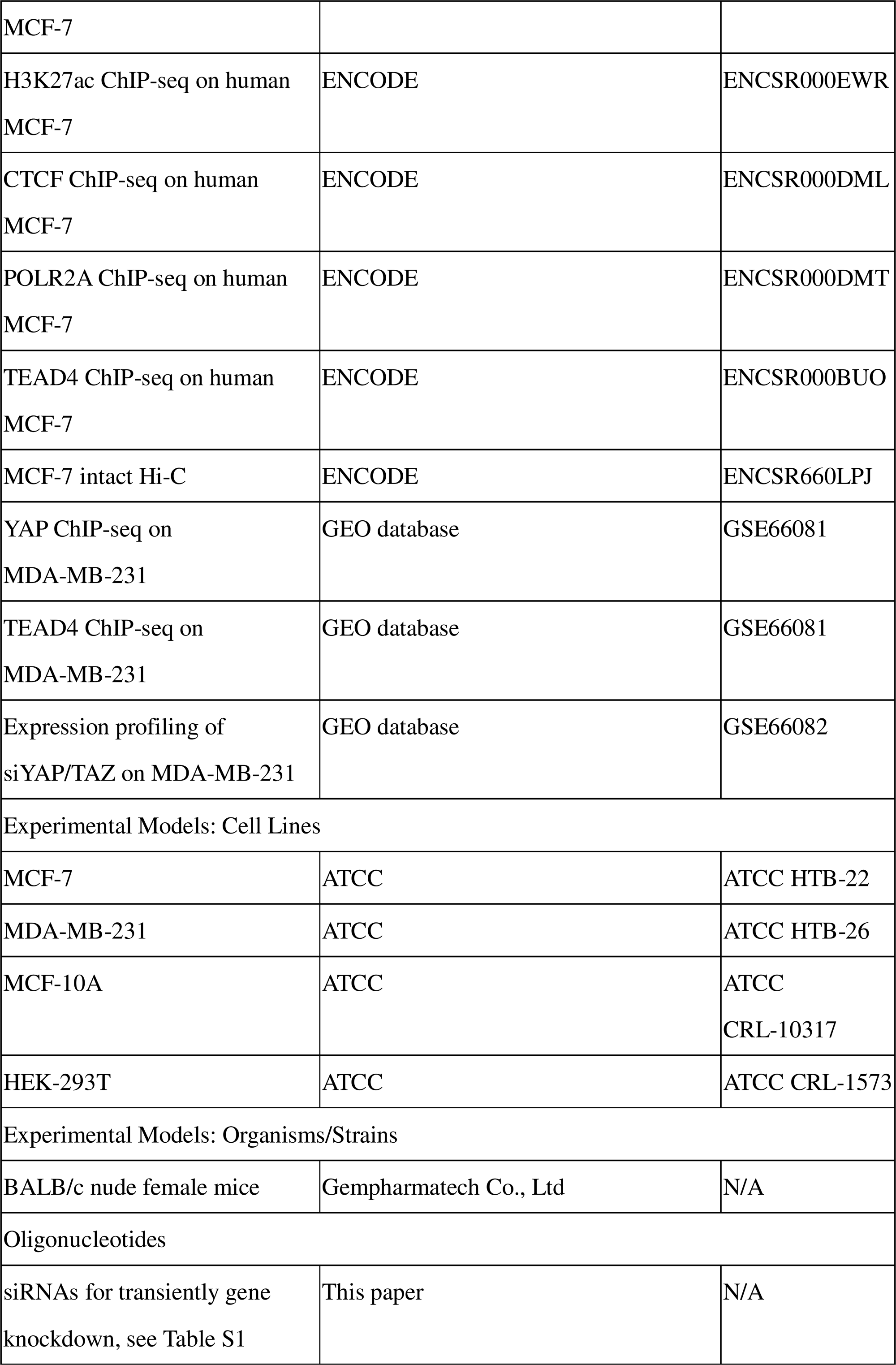
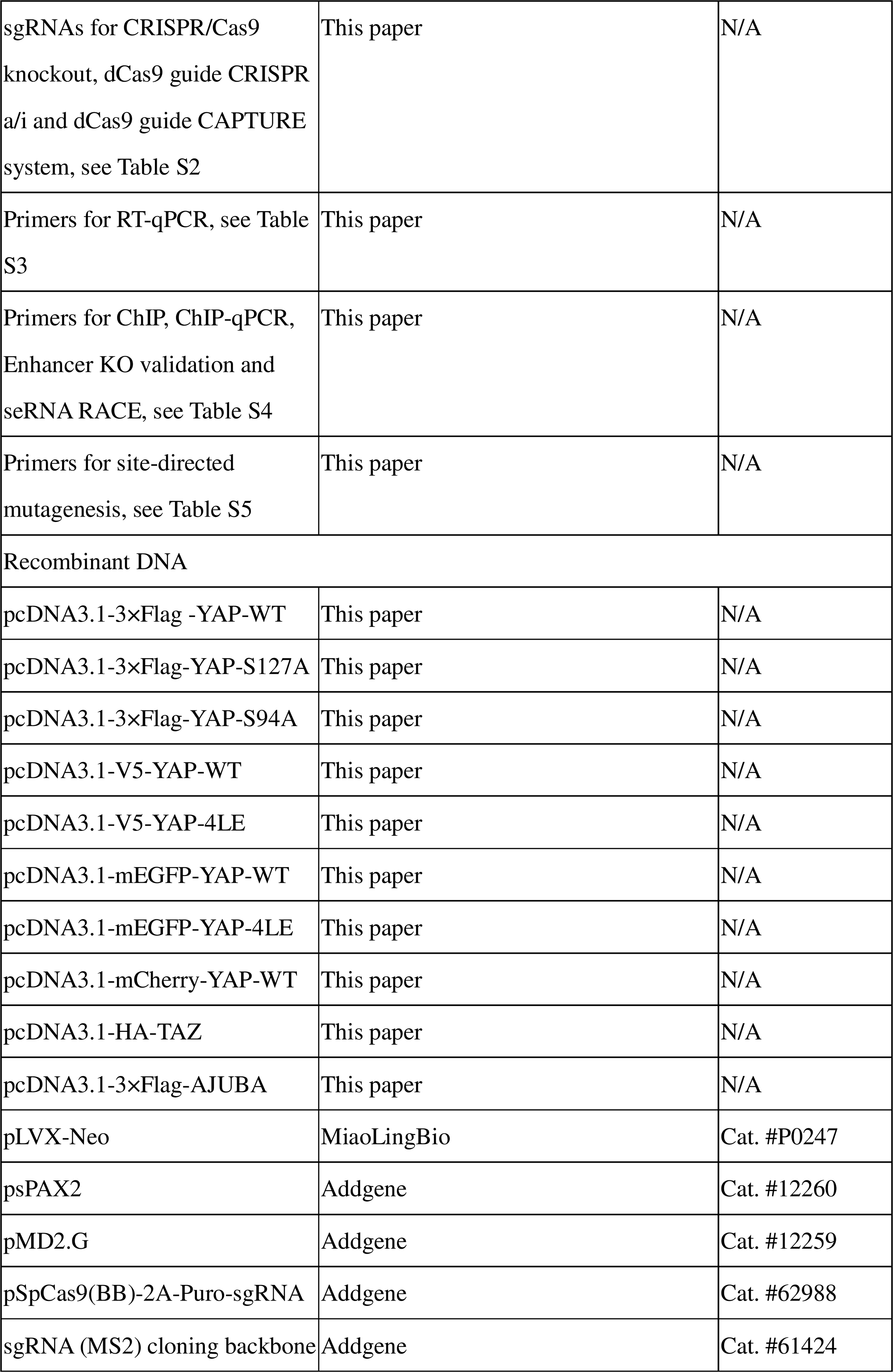
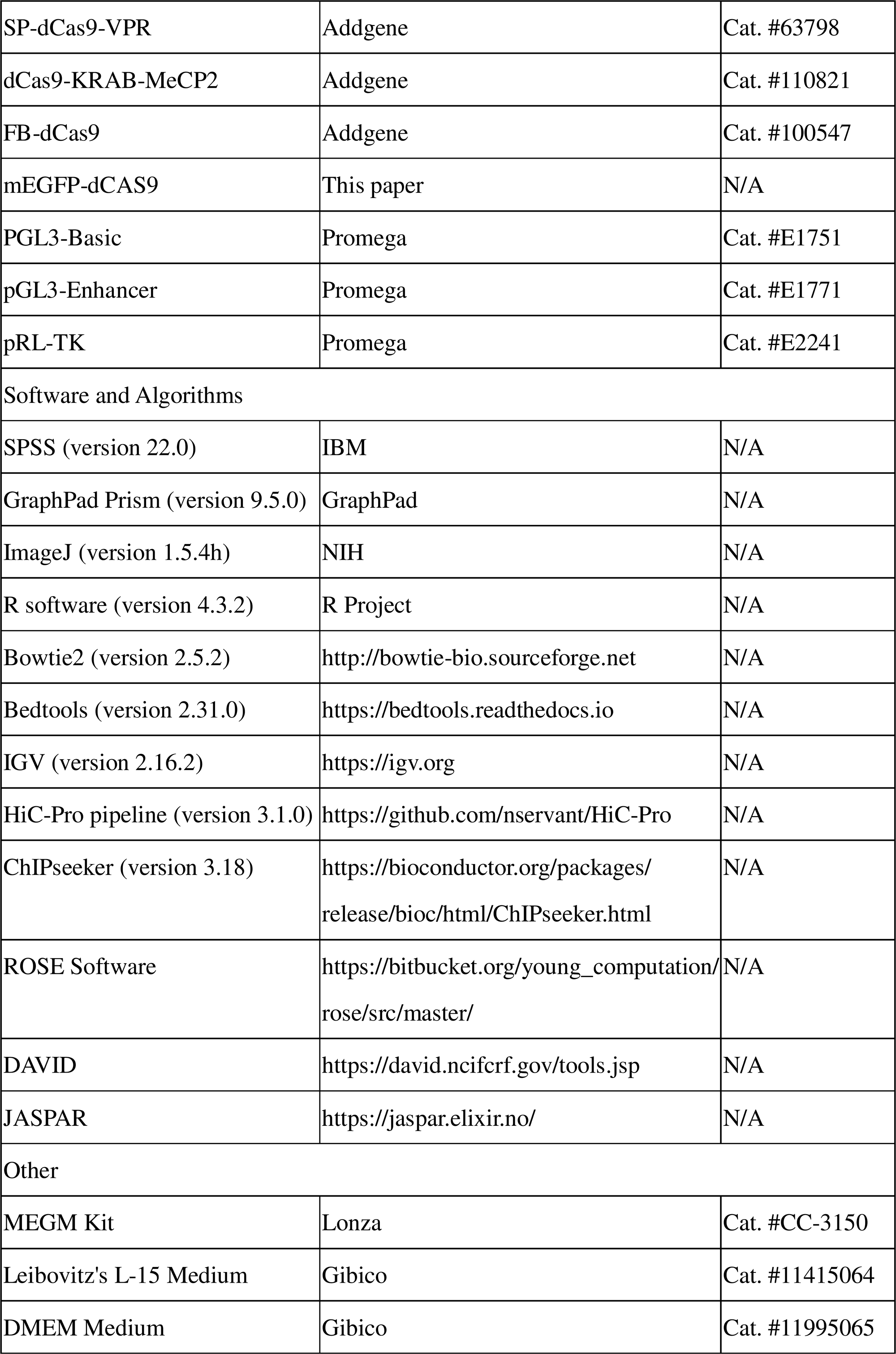
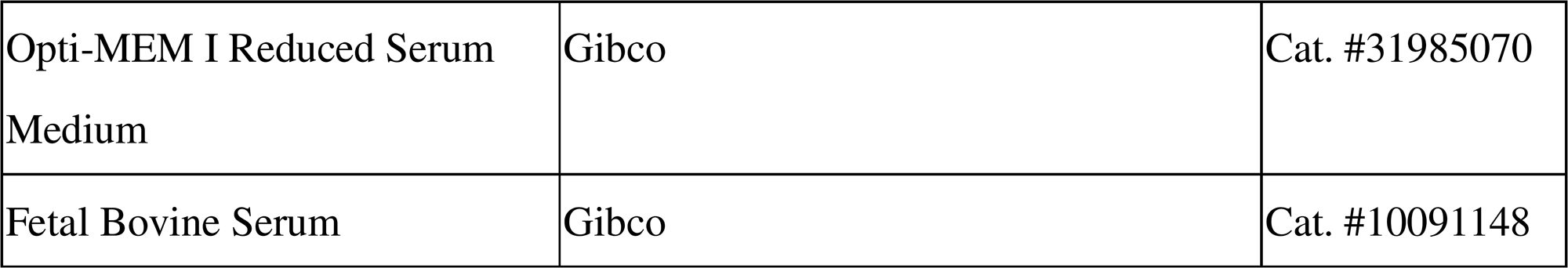

### RESOURCE AVAILABILITY

#### Lead Contact

Further information and requests for resources should be directed to and will be fulfilled by the Lead Contact, Jie Shen and Daxing Xie.

#### Materials Availability

This study did not generate new unique reagents.

#### Data and Code Availability

- All data supporting the findings of this study are available within the main manuscript and the supplementary files.
- This paper does not report original code.
- Any additional information required to reanalyze the data reported in this paper is available from the lead contact upon request.

### EXPERIMENTAL MODEL AND SUBJECT DETAILS

#### Patient samples

A human breast cancer tissue arrays containing primary tumor and paired paracancerous tissue specimens from 75 cases of breast cancer was obtained from Wuhan Baiqiandu Biotech Company (Cat. #BRC1601). Immunochemistry staining of YAP (Cell signaling technology, Cat. #14074) and AJUBA (Abcam, Cat. #ab244285) was performed to validate the expression correlation between the two proteins in multiple tumors.

A human multiple tumor tissue array containing primary tumor specimens from 37 cases of breast cancer, 39 cases of lung cancer, 38 cases of colorectal cancer, 38 cases of prostate cancer and 38 cases of pancreas cancer was purchased from Xian Alenabio Biotech Company (Cat. #BC000119a). Immunochemistry staining was performed as described above to validate the expression correlation between the two proteins in multiple tumors.

#### Animal models

In this study, all 6-week-old female BALB/c nude mice were obtained from Gempharmatech Co., Ltd (Jiangsu, China) and kept in SPF animal facilities. Animal study was performed under the protocols and guidelines approved by the Institutional Animal Care and Use Committee of Huazhong University of Science and Technology, Wuhan, China. Briefly, mice were randomly assigned into different groups. Then, 1 × 10^6^ MDA-MB-231 cells or MCF-7 cells that were managed as indicated were collected and resuspended in 50 μl PBS with 50% Matrigel, and then orthotopically injected into the fourth mammary fat pads of mice. Tumor size was measured every four days, and four weeks after tumor implantation, the mice were sacrificed and the tumor were collected for further experiments.

#### Cell lines and cell culture

MCF-7 and HEK-293T cells were cultured in Dulbecco’s modified Eagle medium (DMEM), MDA-MB-231 cells were cultured in Leibovitz’s L-15 medium (L-15). The medium above was supplemented with 10% fatal bovine serum and 1% penicillin/streptomycin. MCF-10A cells were cultured using the MEGM kit. All cells were cultured in a 5% CO_2_ incubator at 37 □, with the exception of MDA-MB-231 cells which were cultured in 100% air. And all the cell lines were purchased from the ATCC. The drugs Verteporfin, TED-347, MLN8237, and JQ-1 were dissolved in DMSO and prepared as a mother liquor at concentrations of 1 mM (Verteporfin), 10 mM (TED-347), 10 mM (MLN8237), 2 mM (JQ-1), respectively. Cells were treated with the specified drug at the appropriate working concentration and duration, with DMSO utilized as a control at an equivalent volume.

### METHOD DETAILS

#### Plasmid construction, stable cells establishment and transfection

Plasmids were constructed by ligating the DNA fragments and indicated vector after digesting. Gene transient overexpression was conducted in pcDNA3.1 vector, stable overexpression of YAP was conducted in pLVX-Neo vector. For Dual-luciferase reporter assay, cis-regulatory element sequences from E1/E2/promoter or indicated combination of E1/E2/promoter were cloned into PGL3-Basic (promoter, E1+P, E2+P, E1+E2+P) or pGL3-Enhancer (E1, E2, E1+E2), separately. For CRISPR/Cas9 knockout, indicated sgRNAs were cloned into pSpCas9(BB)-2A-Puro-sgRNA vector, and for dCas9 guide CRISPR a/i, sgRNA (MS2) cloning backbone was used to carry sgRNAs. For dCas9 guide CAPTURE system, multiple gRNAs were constructed into sgRNA (MS2) cloning backbone and expressed in tandem from an artificial polycistronic-tRNA-gRNA (PTG) gene, as previously described^54^. The targeted sequences used are listed in Table S1 and S2. Site-directed mutation was constructed using the MultiS Fast Mutagenesis Kit according to the manufacturer’s instructions, and the primers used are listed in Table S5.

Plasmids were transfected using TurboFect, while small interfering RNAs (siRNAs) were transfected using Lipofectamine 2000. siRNAs were obtained from RiboBio Co., Ltd, Guangzhou, China. siRNAs sequences are listed in Table S1, and plasmid information was listed in the KEY RESOURCES TABLE-Recombinant DNA.

Stable YAP overexpressing and control MCF-7 cells lines were generated using a lentiviral system (pLVX, psPAX2, and pMD2.G). Lentiviruses were packaged in HEK-293T cells, then filtered by 0.45□μM filter and concentrated using Universal Virus Concentration Kit. MCF-7 cells were infected and selected using 700 µg/ml of G418 for two weeks. The CRISPR/Cas9 system was used to generate E1KO/E2KO/YAP-E1KO/YAP-E2KO MCF-7 cell lines, and YAP-KO MDA-MB-231 cells lines. Briefly, sgRNAs were designed using the website software E-CRISP (http://www.e-crisp.org/E-CRISP/), and cloned into the pSpCas9(BB)-2A-Puro (PX459) V2.0 plasmid. Engineered plasmids were transfected into indicated cells. 48h after transfection, the cells were selected under 2 μg/mL puromycin (for MCF-7 cells) or 4 μg/mL (for MDA-MB-231 cells), until the cells in the control group died. Surviving cells were seeded in 96-well plates to obtain single-cell clones, and the amplified clones were collected for Sanger sequencing. All stable cell lines were verified by western blotting and qPCR.

#### Quantitative real-time PCR (qPCR)

qPCR was performed as described previously. Briefly, Total RNA was extracted using the TRIzol reagent and cDNA was synthesized using the RT reagent Kit with gDNA Eraser according to the manufacturer’s instructions. qPCR was performed using the TB Green Premix Ex Taq II FAST qPCR Kit in a QuantStudio 3 Real-Time PCR Instrument (Applied Biosystems). The primers used are listed in Supplementary Table S3. Each experiment was performed in triplicate.

#### Immunoblot assays

Immunoblot assays were performed as previously described. Briefly, Total protein was extracted using NP40 lysis buffer supplemented with Protease Inhibitor Cocktail, Phosphatase Inhibitor Cocktail I and Phosphatase Inhibitor Cocktail II. After centrifugation, the supernatant was collected, and protein concentration was measured using the BCA Protein Assay Kits. Samples were diluted using a 5x protein loading buffer [250 mM tris-HCl (pH 6.8), 10% SDS, 30% glycerol, 5% β-mercaptoethanol, and bromophenol blue], and then boiled at 95-100□ for 5 to 10 min. Samples were loading in 10% SDS-PAGE and separated by electrophoresis, then transferred onto PVDF membranes. Membranes were blocked in 5% non-fat milk for 2 h at room temperature, then incubated overnight at 4 □ with the primary antibodies (see the KEY RESOURCES TABLE-antibodies) at a recommended dilution ratio according to the manufacturer’s protocol. After washing, the membranes then were incubated in 1:5000 HRP-conjugated Goat Anti-Rabbit IgG at room temperature for 2 h. Finally, the membranes were visualized using the West Pico PLUS Chemiluminescent Substrate Kit.

#### ChIP

Chromatin immunoprecipitation (ChIP) was performed using Simple ChIP Enzymatic Chromatin IP Kit according the manufacturer’s instructions. The (Polymerase Chain Reaction) PCR was performed using Phanta Max Super-Fidelity DNA Polymerase by the protocol, and the products were analyzed by electrophoresis in a 1% tris-acetate-EDTA (TAE)/ethidium bromide agarose gel. For ChIP-qPCR experiments, ChIP-enriched DNA fragments were quantified using TB Green Premix Ex Taq II FAST qPCR Kit as previously described. The primers are provided in Table S4. Each experiment was performed in triplicate.

#### RACE assay

Rapid amplification of cDNA ends (RACE) was performed using HiScript-TS 5’/3’ RACE Kit. Briefly, total RNA was isolated using TRIZol, and first-strand cDNA was synthesized according to the manufacturer guidelines. Subsequently, the products were used for 5′-RACE and 3′-RACE respectively, and the amplified fragments were analyzed by electrophoresis in a 1% tris-acetate-EDTA (TAE)/ethidium bromide agarose gel. The corresponding gel fragments were purified using Fast Pure Gel DNA Extraction Mini Kit, and sequenced. The gene specific 3′ RACE and 5′ RACE primers are listed in Supplementary Table S4

#### Immunofluorescence

For Immunofluorescence (IF), indicated cells (5 × 10^4^) were seeded on prepared climbing slices in a 24-well plate and cultured in a 37 □ incubator for 16 h. Cells were then washed and fixed in 4% paraformaldehyde for 10 min at 4 □ followed by iced methanol for 10□min and preceded to blocking without permeabilization. Coverslips were incubated with Rabbit anti-γ-Tubulin (1:100) overnight at 4 □, followed by incubating with Dylight 488 Goat Anti-Rabbit IgG (1:200) at room temperature for 2h. And after wash, coverslips were then incubated with Alexa Fluor 555 Rabbit anti-β-Tubulin (1:200) at room temperature for 2h. DAPI was used for the staining of nuclei. The prepared samples were observed using inverted fluorescence microscope (Ts2R-FL, Nikon) at 40× objective. Cells undergoing mitosis with more than two spindle poles were identified as multipolar division. The aberrant mitosis rate was calculated as the percentage of multipolar mitoses out of all mitotic cells (50-80 mitotic cells were counted per sample). Each experiment was performed in triplicate.

#### Chromosome metaphase spreading assay

Indicated cells were treated with colchicine (0.1ug/ml, 37 □, 3 h), digested and collected, then resuspended and incubated in KCl (0.075 M, 37 □, 20 min). Subsequently, cells were fixed in freshly prepared methanol-acetic acid (3:1 vol/vol) and incubated for 30 min at 37 □. After centrifugation, cells were resuspended in a small volume of fixative solution, then dropped onto pre-cold slides and air dried. Slides were stained using Giemsa stain kit, and the chromosome number was analyzed using microscopy (BX53, OLYMPUS) at 100× oil-immersion objective. Aneuploidy is characterized by an abnormal chromosome count per cell, either exceeding or falling below the typical 46 chromosomes in MCF-10A cells, or exhibiting a greater degree of chromosomal number variation in tumour cells.

Percentage of metaphase cells with aberrant chromosomal number were quantified (80-100 metaphase cells were counted per sample). Each experiment was performed in triplicate.

#### H&E staining assay

For H&E staining, paraffin-embedded Xenograft tumor tissue sample slices were routinely dewaxed, rehydrated, and stained using Hematoxylin and Eosin Staining Kit according to the manufacture’s instruction. The results were analysed using microscopy (BX53, OLYMPUS) at 100× oil-immersion objective. Percentage of mitotic cells with aberrant mitosis were quantified (50-80 mitotic cells were counted per sample).

#### Immunohistochemistry

Immunohistochemistry (IHC) staining of YAP and AJUBA in human breast cancer tissue arrays were performed as the manufacture’s instruction. Briefly, slides were dewaxed, rehydrated, and heated in sodium citrate buffer (0.01 M, pH 6.0) for antigen retrieval. Subsequently, endogenous peroxidase was inhibited with 3% hydrogen peroxide and 0.1% sodium aside for 30 min, and nonspecific staining was blocked by 5% bovine serum albumin for 2 h. The slides were subsequently incubated with 1:200 diluted YAP antibody and 1:100 diluted AJUBA antibody at 4 □ overnight and followed by incubating with biotinylated secondary antibodies at room temperature for 2 min. The slides were stained using Immunohistochemistry Kit for Rabbit Primary Antibody, and counterstained with hematoxylin. The results were analyzed using Olympus microscope (BX53, OLYMPUS).

The immunohistochemically stained tissue arrays were scored separately by two experienced pathologists. The expression of YAP and AJUBA was evaluated using the IHC score calculated by multiplying the proportion and intensity score. The proportion score represents the proportion of positively stained cells: 0 (<5%), 1 (5-25%), 2 (26-50%), 3 (51-75%), and 4 (>75%), and the intensity score reflects the staining intensity (0, no staining; 1, weak; 2, moderate; 3, strong). An IHC score of ≤5 was assessed as low expression and scores of 6–12 was evaluated as high.

#### Dual-luciferase reporter assay

Dual-luciferase reporter assay was performed using the Dual-Luciferase Reporter Assay Kit. Briefly, pGL3-Basic/Enhancer plasmids with inserted target sequences, pRL-TK plasmid and indicated YAP plasmids were co-transfected into HEK-293T cells. After 48 h, cells were lysed to collect the supernatant, and the firefly luciferase activity was assayed and normalized to that of Renilla luciferase. Each experiment was performed in triplicate.

#### In Situ CAPTURE

In Situ CAPTURE (CRISPR Affinity Purification in situ of Regulatory Elements) was performed as described^30^. Briefly, cells (5×10^7^), which transfected with FB-dCas9 plasmid and AJUBA super-enhancer E1/E2 sgRNAs or non-targeting sgRNA constructed plasmids, were cross-linked with 1% formaldehyde for 10 min, and quenched with 0.25 M glycine for 5 min. Cells were lysed and centrifuged to isolate the nuclei, and then were resuspended and sonicated into 200∼500 bp length of segments. Supernatant was then incubated with Streptavidin T1 Dynabeads at 4 □ overnight and followed by wash with low-salt buffer and high-salt buffer. For obtention of dCas9-captured DNA, the chromatin fragments were eluted, reverse cross-linked, and purified using Simple ChIP Enzymatic Chromatin IP Kit. The products were amplified and analyzed by electrophoresis in a 1% tris-acetate-EDTA (TAE)/ethidium bromide agarose gel or subjected to qPCR to detect the captured promoter of AJUBA. For obtention of dCas9-captured proteins, the Streptavidin T1 Dynabeads were washed with IP binding buffer and suspended in 1x protein loading buffer, then incubated at 95-100 □ for 20 min. The proteins were separated by SDS-PAGE and analyzed by Western blot. The sgRNA sequences and primers used are listed in Table S2 and S4. Each experiment was performed in triplicate.

#### CRISPR a/i

CRISPR activation/interference (CRISPR a/i) was performed as described. Briefly, sgRNAs targeting AJUBA super-enhancer E1/E2 were cloned into sgRNA (MS2) cloning backbone plasmid. For CRISPR activation or interference, SP-dCas9-VPR or dCas9-KRAB-MeCP2 were co-transfected with constructed sgRNA plasmids into MCF-7 or MDA-MB-231 cells, respectively. After 48 h, the cells were harvested for western blot or qPCR analysis. The sgRNA sequences are listed in Table S2. Each experiment was performed in triplicate.

#### Live cell imaging and FRAP

For live cell imaging, 4 × 10^4^ MCF-7 cells were seeded on a 24-well glass-bottom confocal plate and transfected with the mEGFP-YAP plasmid. For 1,6-hexanediol treatment, indicated cells were treated with PBS or 3% 1,6-hexanediol, and observe the real-time status change of mEGFP-YAP phase-separation condensates. Images were acquired on confocal microscope (FV3000, Olympus) at 100× oil-immersion objective with cell culture system.

For fluorescence recovery after photobleaching (FRAP), on the basis of live cell imaging, one of YAP phase-separation condensates was identified and drawn a region of interest (ROI) of it by using the Olympus FV3000 imaging software within the stimulation module. Then the ROI was subjected to photo-bleach by utilizing the 488 nm laser line at 20% laser power for 250 ms, and images were collected every 1 s post-bleaching. Fluorescence intensity was measured using the ImageJ. FRAP experiment was performed in triplicate.

For colocalization analysis, cells were prepared as previously described, and co-transfected with the mCherry-YAP plasmid, mEGFP-dCas9 plasmid and constructed sgRNA plasmids targeting AJUBA super-enhancer E1/E2. Cells were imaged on confocal microscope (FV3000, Olympus) at 100× oil-immersion objective.

#### RNA-FISH

RNA fluorescence in situ hybridization (RNA-FISH) was performed using Fluorescent In Situ Hybridization Kit according to the manufacture’s protocol. Briefly, indicated cells were grown on climbing slices in a 24-well plate, then fixed in 4% paraformaldehyde at room temperature for 10 min. After permeabilized, cells were incubated with pre-hybridization buffer for 30 min at 37 □, and then incubated with pre-heated hybridization buffer with FISH probe mix added at 37 □ overnight, protected from light. Subsequently, climbing slices were washed and stained with DAPI, and then sealed with Antifade Mounting Medium. Images were collected using inverted fluorescence microscope (Ts2R-FL, Nikon) at 60× objective. FISH probes targeting AJUBA seRNA were designed and synthesized by Guangzhou RiboBio Co., LTD. Each experiment was performed in triplicate.

#### ChIP-seq and Hi-C data analysis

Chromatin immunoprecipitation sequencing (ChIP-seq) data of H3K4me1, H3K4me3, H3K27ac, CTCF, POLR2A, and TEAD4 on MCF-7 cells were obtained from the ENCODE database (https://www.encodeproject. org/). The Rank Ordering of Super Enhancers (ROSE) algorithm was used to identify SEs from H3K27ac ChIP-seq data^55^. Alignment of the sequencing data to the human reference genome (GRCh38) was performed using Bowtie2 and Bedtools, and the bigwig format files of ChIP-seq date above were visualized using the Integrative Genomics Viewer (IGV). High throughput chromosome conformation capture (Hi-C) raw date of MCF-7 was also obtained from the ENCODE database. Quality control procedures were applied to the raw data, involving the removal of low-quality reads and appropriate trimming to ensure the data quality. And then, clean reads were processed the HiC-Pro pipeline according to the instruction^56^. Subsequently, reads were mapped to the human reference genome (GRCh38), and the identified chromatin loops were visualized on the WashU EpiGenome Browser (http://epigenomegateway.wustl.edu). And date obtained from ENCODE database were provided in the KEY RESOURCES TABLE-Deposited Data.

#### Bioinformatic analysis

ChIP-seq data of YAP and TEAD4 on MDA-MB-231 were obtained from GEO database (GSE66081) and annotated using ChIPseeker tool. Meanwhile, the expression profile of siYAP/TAZ v.s. siNC on MDA-MB-231 cell was download from GEO database (GSE66082). The different expression genes with fold change (FC) >= 2 and p-value < 0.01 were intersected with the genes from ChIP-seq data containing both YAP binding peaks and TEAD4 binding peaks. Gene ontology (biological process) enrichment of these intersected genes were performed using DAVID software.

### QUANTIFICATION AND STATISTICAL ANALYSIS

SPSS (version 22.0) and GraphPad Prism (version 9.5.0) were used for statistical analysis. Continuous data are presented as mean ± standard deviation (SD) and statistically analyzed using Student’s t-test (two-tailed) or analysis of variance (ANOVA). Enumeration data were analyzed using the Fisher’s exact test. Survival was analyzed using the Kaplan-Meier curve with a log-rank test. Statistical significance was set as ns, not statistically significant, *p < 0.05, **p < 0.01, ***p < 0.001, and ****p < 0.0001.

