## Supplemental tables and figures for "YAP promotes chromosomal instability by transcriptionally activating AJUBA through its super enhancer in breast cancer"

*SUPPLEMENTARY INFORMATIONS*

**Supplementary figures**

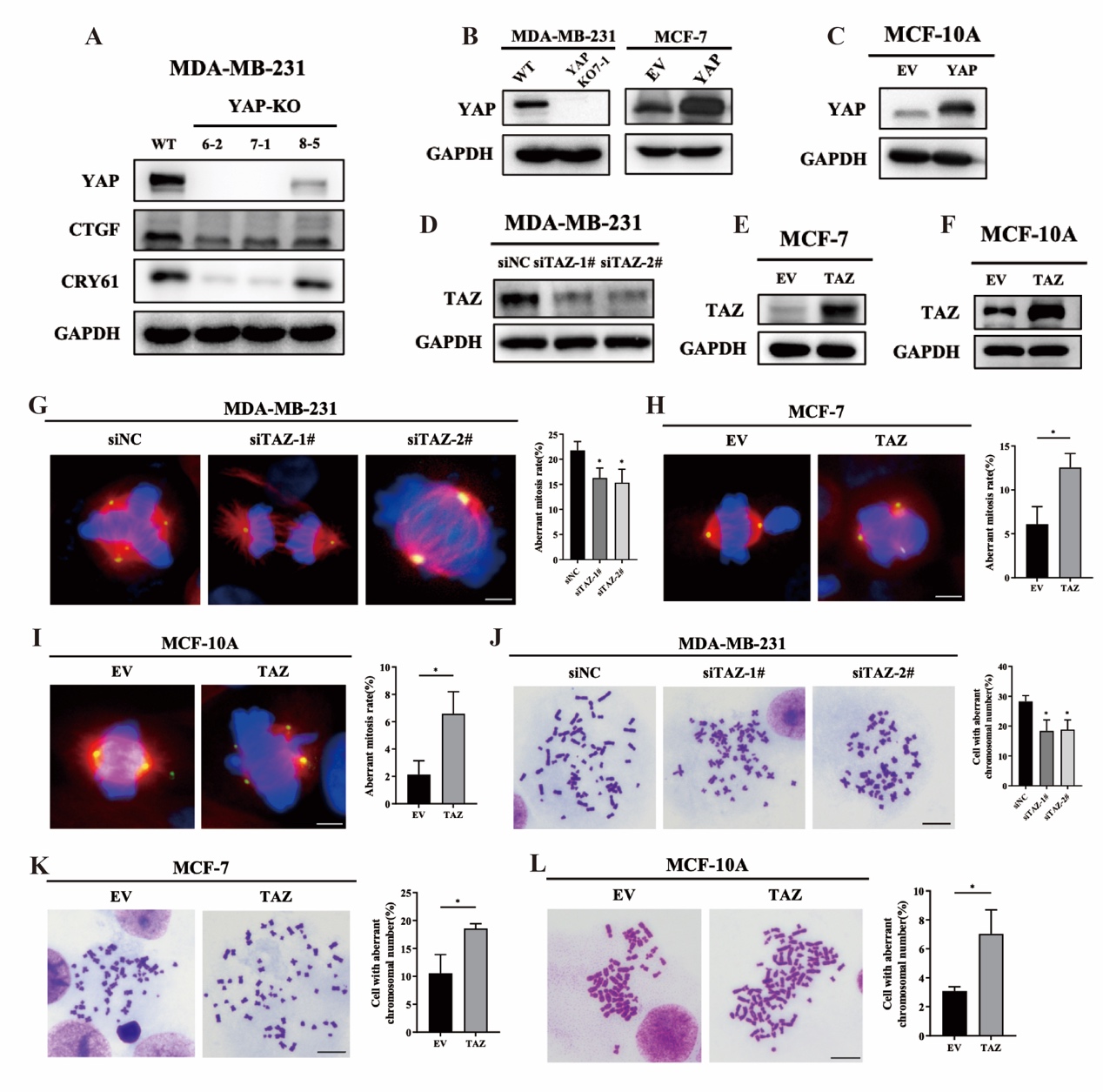

**Figure S1. YAP induces CIN formation and aberrant mitosis in breast cancer.**

(A) Cell lysates from MDA-MB-231 wild type (WT) and three YAP-knockout clones (KO-1, KO-2, KO-3) were used for immunoblot and probed for YAP, CTGF, CYR61. GAPDH was used as a loading control.

(B, C) Cell lysates from MDA-MB-231 with wild type (WT) or YAP knockout (YAP-KO), MCF7 with empty vector (EV) or YAP overexpression (YAP) (B), and MCF-10A with empty vector (EV) or YAP overexpression (YAP) (C) were used for immunoblot. Lysates were probed for YAP and GAPDH was used as a loading control.

(D, E and F) Cell lysates from MDA-MB-231 cells transfected with scramble (siNC) or TAZ interfering siRNA (siTAZ-1# and siTAZ-2#) (D), MCF7 with empty vector (EV) or TAZ overexpression (TAZ) (E), and MCF-10A with empty vector (EV) or TAZ overexpression (YAP) (F) cells were used for immunoblot. Lysates were probed for TAZ and GAPDH was used as a loading control.

(G, H and I) Representative images of aberrant mitosis from MDA-MB-231 cells transfected with scramble (siNC) or TAZ interfering siRNA (siTAZ-1# and siTAZ-2#) (G), MCF7 with empty vector (EV) or TAZ overexpression (H), and MCF-10A with empty vector (EV) or TAZ overexpression (YAP) (I). β-Tubulin (red), γ-Tubulin (green), Nuclei (blue). Scale bar: 10μm. Histograms show the mean percentage ± SD of aberrant mitosis rate. *p<0.05; **p<0.01.

(J, K and L) Representative images of chromosome metaphase spreading from MDA-MB-231 cells transfected with scramble (siNC) or TAZ interfering siRNA (siTAZ-1# and siTAZ-2#) (J), MCF7 with empty vector (EV) or TAZ overexpression (K), and MCF-10A with empty vector (EV) or TAZ overexpression (YAP) (L). Scale bar: 5μm. Histograms show the mean percentage ± SD of cell rate with aberrant chromosomal number. *p<0.05; ***p<0.001.

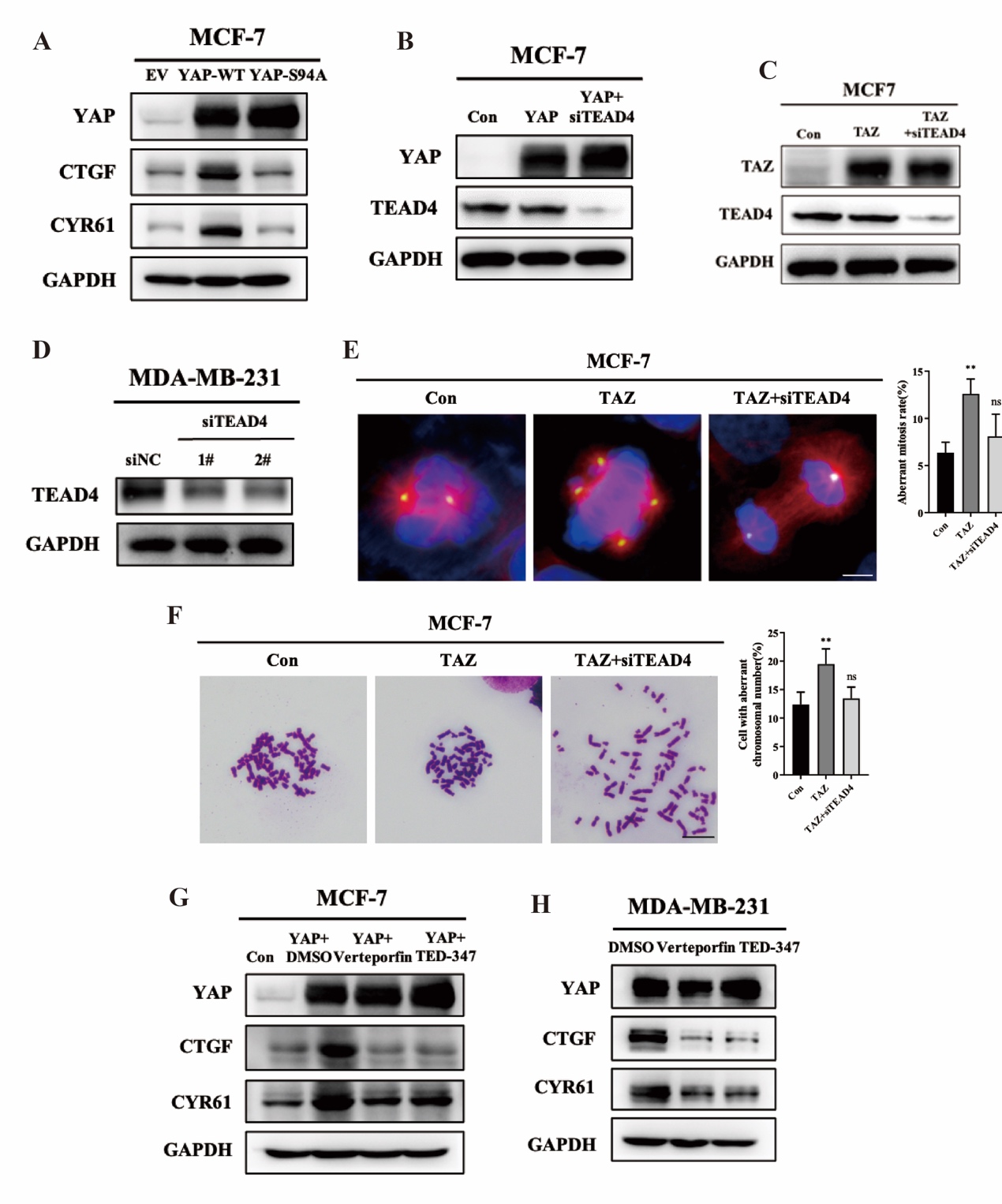

**Figure S2. YAP-TEAD interaction is essential in CIN formation and aberrant mitosis.**

(A) MCF-7 cells transfected with empty vector (EV), wild type YAP (YAP WT) and YAP-S94A mutant (YAP S94A) were used for immunoblot. Cell lysates were probed for YAP, CYR61 and CTGF. GAPDH was used as a loading control.

(B) MCF-7 cells transfected with wild type YAP plasmid and/or siTEAD4 siRNA were used for immunoblot. Cell lysates were probed for YAP and TEAD4 Empty vector and scramble siRNA were used as negative control, and GAPDH was used as a loading control.

(C) MCF-7 cells transfected with siTEAD4 siRNAs or TAZ overexpression plasmids were collected for immunoblot. Empty vector and scramble siRNA were used as negative control. Cell lysates were probed for TAZ and TEAD4. GAPDH was used as a loading control.

(D) MDA-MB-231 cells transfected with scramble (siNC) or siTEAD4 siRNAs (siTEAD4-1#, siTEAD4-2#) were used for immunoblot. Cell lysates were probed for TEAD4. GAPDH was used as a loading control.

(E) Representative images of aberrant mitosis in MCF7 cells transfected with siTEAD4 siRNAs or TAZ overexpression plasmids. Empty vector and scramble siRNA were used as negative control. β-Tubulin (red), γ-Tubulin (green), Nuclei (blue). Scale bar: 10 μm, histograms show the mean percentage ± SD of aberrant mitosis rate. ns., no significant difference; **p<0.01.

(F) Representative images of chromosome metaphase spreading in MCF7 cells transfected with siTEAD4 siRNAs or TAZ overexpression plasmids. Empty vector and scramble siRNA were used as negative control. Scale bar: 5μm, histograms show the mean percentage ± SD of cell rate with aberrant chromosomal number. ns., no significant difference; **p<0.01.

(G) MCF-7 was stablely transfected with control (Con) or YAP overexpressing plasmid. Then cells were treated with DMSO, Verteporfin (1μM) or TED-347 (10μM) for 24 h. Western blot assay was performed to examine the protein level of YAP, CYR61 and CTGF. GAPDH was used as a loading control.

(H) MDA-MB-231 cells were treated with DMSO, Verteporfin (1μM) or TED-347 (10μM) for 24 h. Western blot assay was performed to examine the protein level of YAP, CYR61 and CTGF. GAPDH was used as a loading control.

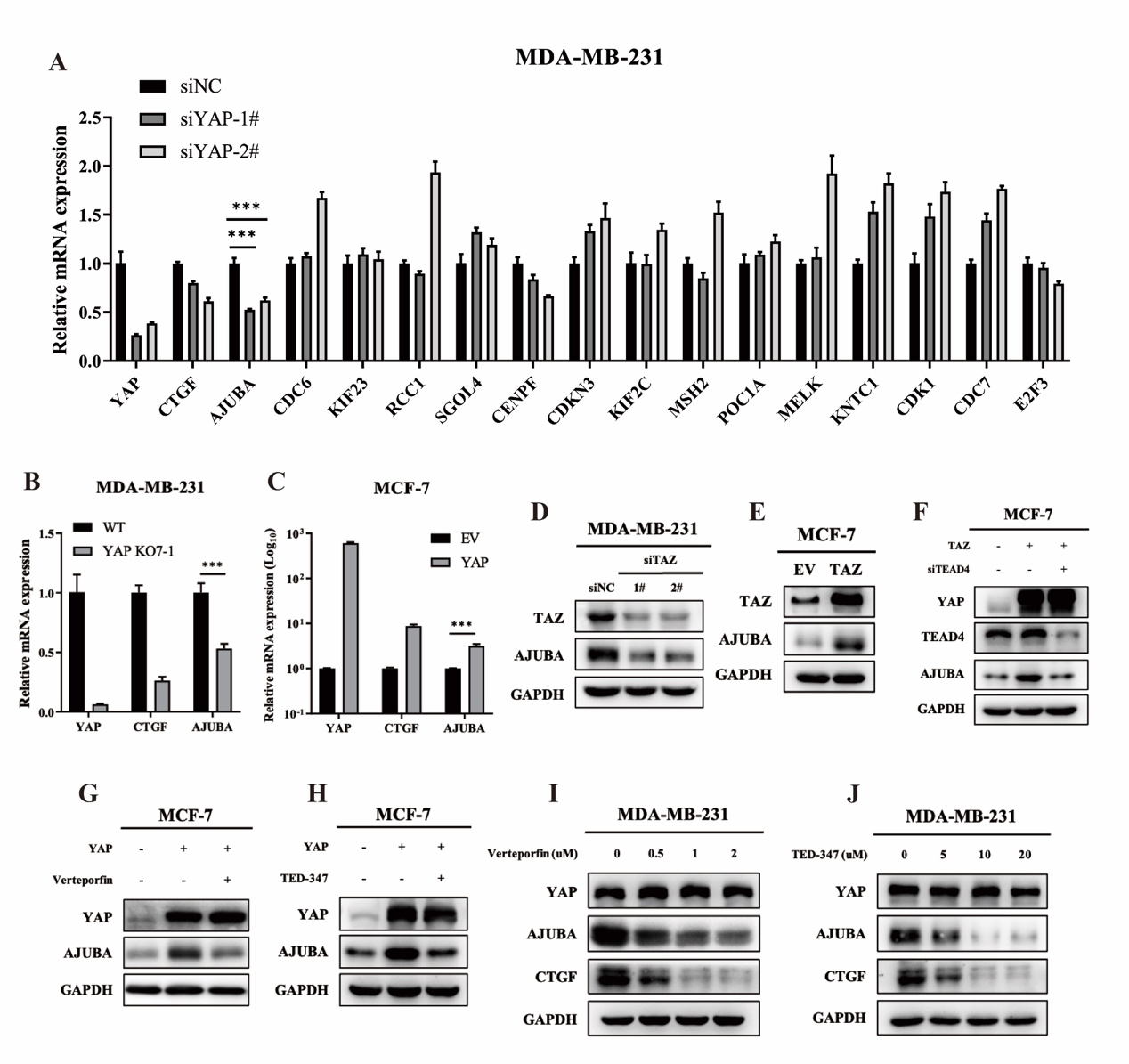

**Figure S3. YAP/TEAD transcriptionally activating spindle assembly checkpoint AJUBA expression.**

(A) The expression level of 15 candidate genes in “Mitotic cell cycle process” category from figure 3C was evaluated in MDA-MB-231 cells with control (siNC) or YAP knockdown (siYAP-1#, siYAP-2#) via qPCR assay. CTGF was the direct transcriptional target of YAP and GAPDH was used as an endogenous control. ***p<0.001.

(B and C) qPCR analysis of AJUBA expression level in MDA-MB-231 cells with wild type (WT) or YAP knockout (YAP KO7-1) (B), and MCF-7 with empty vector (EV) or YAP overexpression (C). GAPDH was used as an endogenous control.

(D) MDA-MB-231 cells were transfected with scramble (siNC) or siTAZ siRNAs (siTAZ-1#, siTAZ-2#). Cell lysates were used for immunoblot, and probed for TAZ and AJUBA. GAPDH was used as a loading control.

(E) MCF7 cells were transfected with empty vector (EV) or TAZ overexpressing plasmid (TAZ). Cell lysates were used for immunoblot, and probed for TAZ and AJUBA. GAPDH was used as a loading control.

(F) MCF-7 cells transfected with siTEAD4 siRNA and/or TAZ overexpressing plasmid were collected for immunoblot. Scramble siRNA and empty vector were used as negative control. Cell lysates were probed for TAZ, AJUBA and TEAD4, GAPDH was used as a loading control.

(G and H) Empty vector (EV) or YAP plasmids were transfected into MCF-7. Then cells were treated with Verteporfin at a dose of 1 μM (G) or TED-347 at a dose of 10 μM (H) for 24 h. DMSO was used as negative control. Cell lysates were collected for immunoblot and probed for YAP and AJUBA. GAPDH was used as a loading control.

(I and J) MDA-MB-231 cells were treated with DMSO or indicated concentration of Verteporfin (I) or TED-347 (J) for 24 h. Cell lysates were collected for immunoblot and probed for YAP, AJUBA and CTGF. GAPDH was used as a loading control.

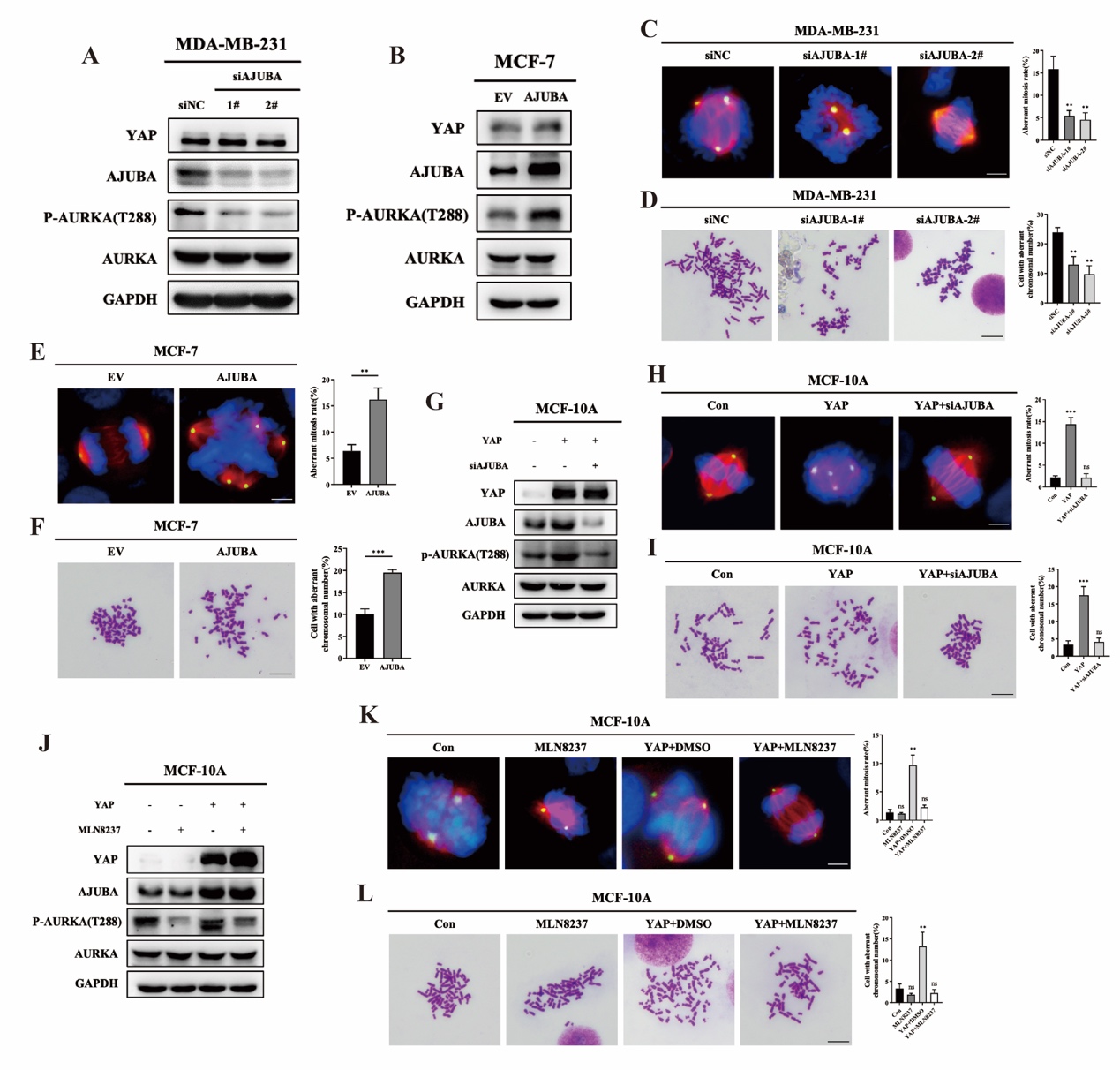

**Figure S4. YAP regulates CIN through AJUBA-AURKA axis.**

(A) MDA-MB-231 cells were transfected with scramble (siNC) or siAJUBA siRNAs (siAJUBA-1#, siAJUBA-2#). Cell lysates were used for immunoblot, and probed for YAP, AJUBA, AURKA, and p-AURKA (T288). GAPDH was used as a loading control.

(B) MCF7 cells were transfected with empty vector (EV) or AJUBA overexpressing plasmid (AJUBA). Cell lysates were used for immunoblot, and probed for YAP, AJUBA, AURKA, and p-AURKA (T288). GAPDH was used as a loading control.

(C) Representative images of aberrant mitosis in MDA-MB-231 cells transfected with siAJUBA siRNAs. Scramble siRNA were used as negative control. β-Tubulin (red), γ-Tubulin (green), Nuclei (blue). Scale bar: 10μm, histograms show the mean percentage ± SD of aberrant mitosis rate. **p<0.01.

(D Representative images of chromosome metaphase spreading in MDA-MB-231 cells transfected with siAJUBA siRNAs. Scramble siRNA were used as negative control. Scale bar: 5μm, histograms show the mean percentage ± SD of cell rate with aberrant chromosomal number. **p<0.01.

(E) Representative images of aberrant mitosis in MCF7 cells transfected with empty vector (EV) or AJUBA overexpressing plasmid (AJUBA). β-Tubulin (red), γ-Tubulin (green), Nuclei (blue). Scale bar: 10μm, histograms show the mean percentage ± SD of aberrant mitosis rate. **p<0.01.

(F) Representative images of chromosome metaphase spreading in MCF7 cells transfected with empty vector (EV) or AJUBA overexpressing plasmid (AJUBA). Scale bar: 5μm, histograms show the mean percentage ± SD of cell rate with aberrant chromosomal number. ***p<0.001.

(G) MCF-10A cells were transfected with YAP overexpressing plasmid and/or siAJUBA siRNAs (siAJUBA-1#, siAJUBA-2#). Empty vector and scramble siRNA were used as negative control. Cell lysates were probed for YAP, AJUBA, AURKA, and p-AURKA (T288). GAPDH was used as a loading control.

(H) Representative images of aberrant mitosis in MCF-10A cells transfected with YAP overexpressing plasmid and/or siAJUBA siRNAs (siAJUBA-1#, siAJUBA-2#). Empty vector and scramble siRNA were used as negative control. β-Tubulin (red), γ-Tubulin (green), Nuclei (blue). Scale bar: 10 μm, histograms show the mean percentage ± SD of aberrant mitosis rate. ns. no significant difference, ***p<0.001.

(I) Representative images of chromosome metaphase spreading in MCF-10A cells transfected with YAP overexpressing plasmid and/or siAJUBA siRNAs (siAJUBA-1#, siAJUBA-2#). Empty vector and scramble siRNA were used as negative control. Scale bar: 5μm, histograms show the mean percentage ± SD of cell rate with aberrant chromosomal number. ns. no significant difference, ***p<0.001.

(J) MCF-10A cells with empty vector (EV) or YAP overexpression were treated with DMSO or MLN8237 at a dose of 10μM for 24 h. Then cells were collected for immunoblot. Cell lysates were probed for YAP, AJUBA, AURKA, and p-AURKA (T288). GAPDH was used as a loading control.

(K) MCF-10A cells with empty vector (EV) or YAP overexpression were treated with DMSO or MLN8237 at a dose of 10μM for 24 h. Then immunofluorescence was performed and representative images of aberrant mitosis were shown. β-Tubulin (red), γ-Tubulin (green), Nuclei (blue). Scale bar: 10μm, histograms show the mean percentage ± SD of aberrant mitosis rate. ns. no significant difference, **p<0.01.

(L) MCF-10A cells with empty vector (EV) or YAP overexpression were treated with DMSO or MLN8237 at a dose of 10μM for 24 h. Then karyotype analysis was performed and representative images of chromosome metaphase spreading were shown. Scale bar: 5μm, histograms show the mean percentage ± SD of cell rate with aberrant chromosomal number. no significant difference, **p<0.01.

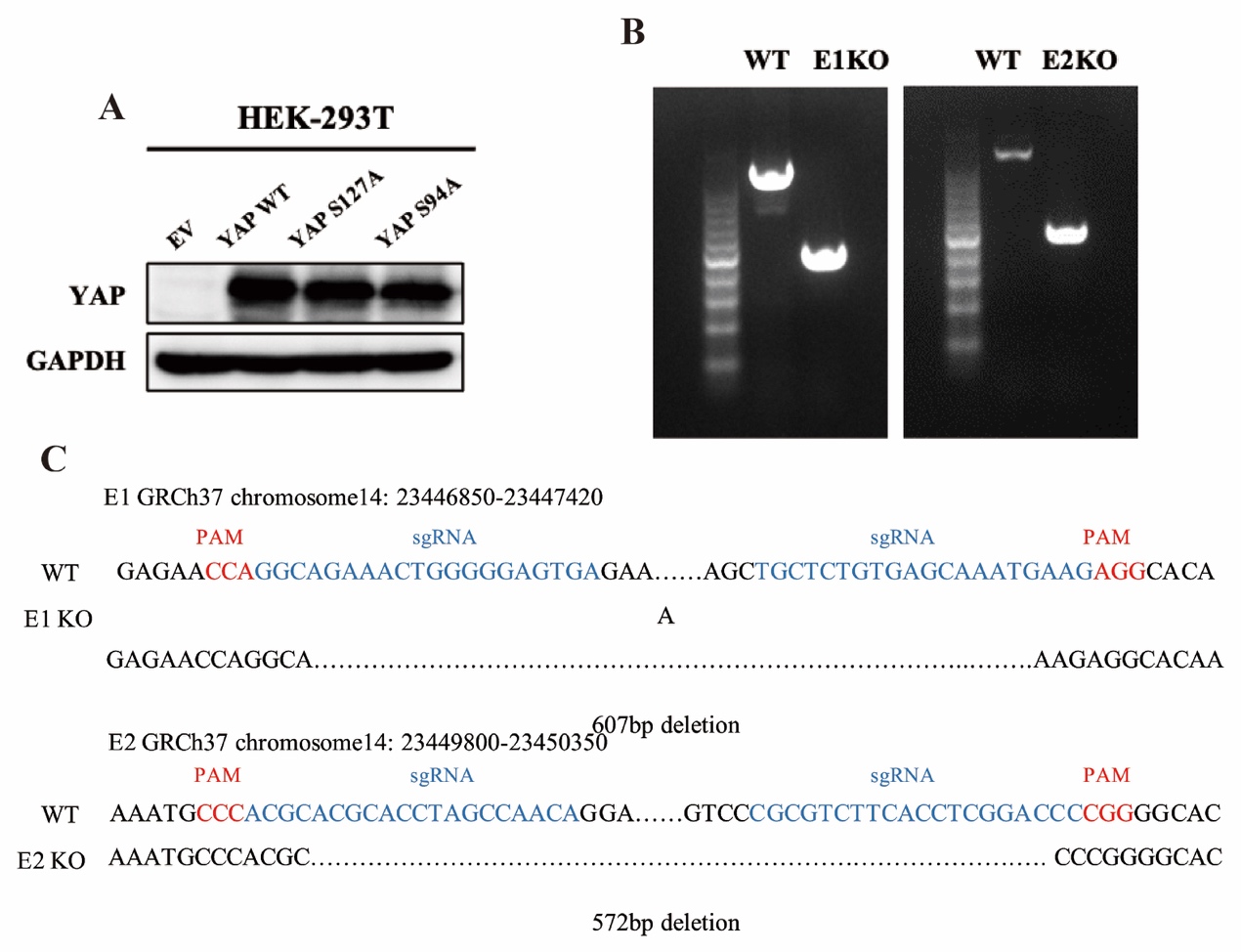

**Figure S5. YAP/TEAD promotes AJUBA transcription through its super enhancer.**

(A) Cell lysates from HEK-293T cells transfected with empty vector (EV), wild type YAP (YAP WT) and YAP-S94A mutant (YAP S94A) were collected for immunoblot. Lysates were probed for YAP and GAPDH was used as a loading control.

(B) PCR amplification and agarose gel electrophoresis were performed to analysis the exact genome DNA from MCF7 cell with wild type (WT) or E1/E2 knockout (E1/E2-KO). Primers were designed within ~250 bp of deleted region.

(C) Sanger sequencing analysis was performed using the PCR products from Fig.S5B to validate the deleted regions on AJUBA enhancer. The PAM region was marked as red and the sgRNA targets were marked as blue.

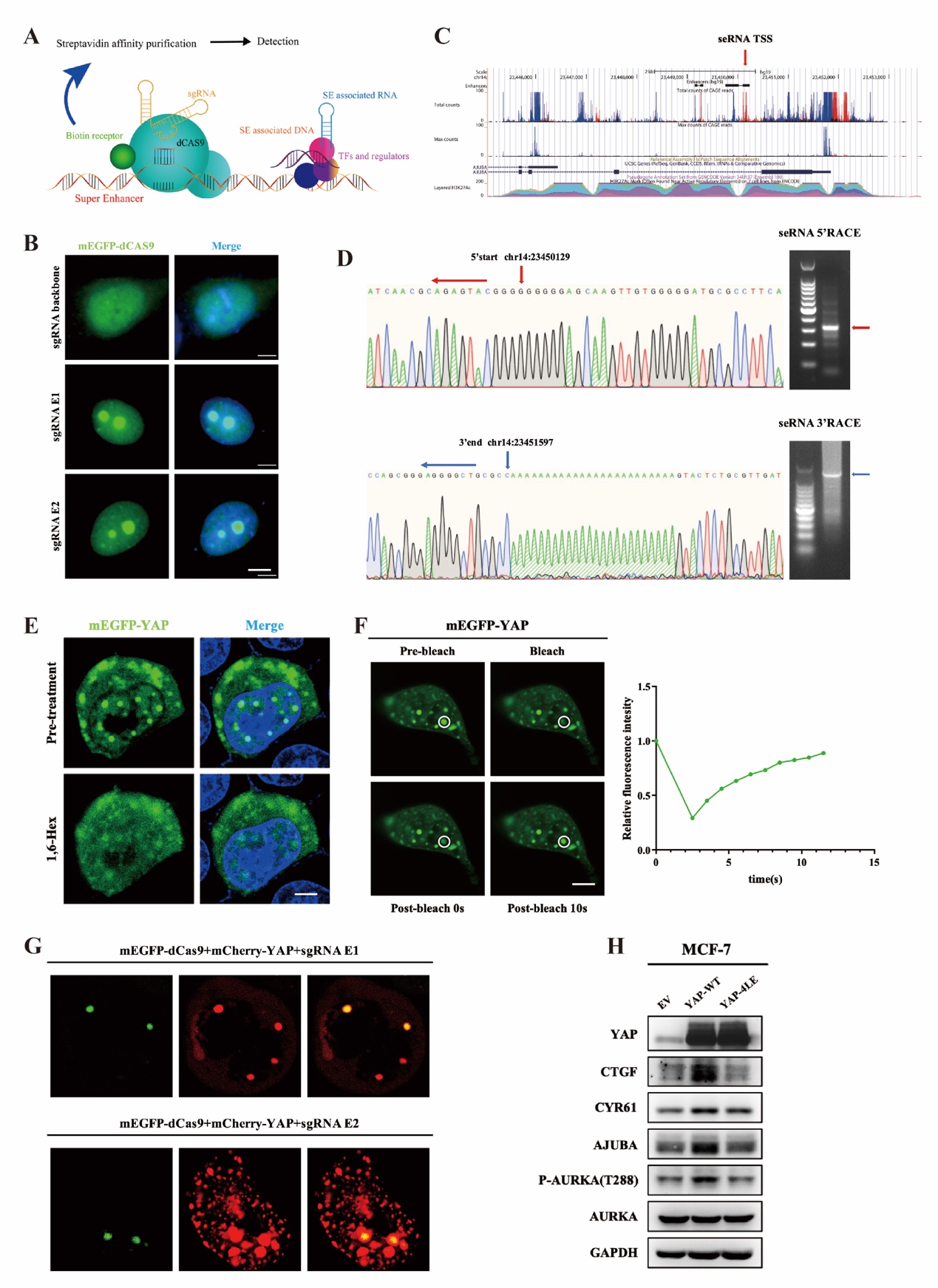

**Figure S6. YAP/TEAD directly activates super enhancer and induces SE-promoter proximity on AJUBA gene.**

(A) Schematic of in situ CAPTURE.

(B) Immunofluorescence was used to evaluate the CAPTURE efficiency on E1/E2 loci of AJUBA gene. E1 and E2 marked by sgRNAs were interacted with mEGFP fused dCAS9 (green). DAPI (blue) was used for nuclei staining. Sale bar: 5μm.

(C) Predicted seRNA transcriptional start site (TSS) of AJUBA according to FANTOM5 database.

(D) RACE assay and sanger sequencing analysis were used to validate AJUBA seRNA sequence. The 5’ start and 3’ end of AJUBA seRNA were pointed out by red arrow and blue arrow, respectively.

(E) MCF-7 cells transfected with mEGFP-YAP (green) were observed for the formation of YAP phase separation under confocal microscopy before and after treatment with PBS or 3% 1,6-Hex for 30 s. Hoechst 33342 (blue) was used for nuclei staining. Scale bar: 5μm.

(F) Fluorescence recovery after photobleaching (FRAP) assay in MCF7 cells transfected with mEGFP-YAP. The bleached and control YAP condensate (white circle) was marked by red and yellow rectangle, respectively (left). Quantifications of relative fluorescence intensity of bleached YAP condensate at indicate time point were shown (right). Data are shown as mean ± SEM. Scale bar: 5μm.

(G) Immunofluorescence showing mCherry-YAP (red) co-localized with E1 and E2 locus of AJUBA gene in MCF-7 cells. E1 and E2 were marked by sgRNA conjugated with mEGFP fused dCAS9 sgRNAs (green). Scale bar: 5μm.

(H) MCF-7 cells transfected with empty vector (EV), wild type YAP (YAP-WT) and YAP-4LE mutant were collected for immunoblot. Cell lysates were probed for YAP, CTGF, CRY61, AJUBA, AURKA, and p-AURKA (T288). GAPDH was used as a loading control.

**Supplementary Table**

| **Table S1. siRNAs for transiently gene knockdown, related to STAR Methods.** | |
| --- | --- |
| Name | Target Sequence (5`-3`) |
| siYAP-1# | GCGTAGCCAGTTACCAACA |
| siYAP-2# | GGTGATACTATCAACCAAA |
| siTAZ-1# | TGCTTCCTCAGTTACACAAAG |
| siTAZ-2# | TCCTAACAGTCCGCCCTACTT |
| siTEAD4-1# | GCTTGTGGATGAAGTTGAT |
| siTEAD4-2# | GGACACTACTCTTACCGCA |
| siAJUBA-1# | TGGACCGGGATTATCACTTTG |
| siAJUBA-2# | GTGTCTGTGGTCACTTGATTT |

**Table S2. sgRNAs for CRISPR/Cas9 knockout, dCas9 guide CRISPR a/i and dCas9 guided CAPTURE system, related to STAR Methods.**

| Name | Target Sequence (5`-3`) |
| --- | --- |
| sgRNAs for YAP KO | |
| sgRNA-YAP-1# | GCAGTCGCATCTGTTGCTGC |
| sgRNA-YAP-2# | GAGCACTCTGACTGATTCTC |
| sgRNA-YAP-3# | ACATCGATCAGACAACAACA |
| sgRNAs for E1/E2 KO | |
| E1KO-1# | TCACTCCCCCAGTTTCTGCCTGG |
| E1KO-2# | TGCTCTGTGAGCAAATGAAGAGG |
| E2KO-1# | TGTTGGCTAGGTGCGTGCGTGGG |
| E2KO-2# | CGCGTCTTCACCTCGGACCCCGG |
| sgRNAs for CRISPR a/i | |
| sgRNA-E1-1# | GTGAAAGGGGGGAGCAAGTT |
| sgRNA-E1-2# | GAATTCGGAGAATGCGCGGG |
| sgRNA-E2-1# | GTGCGCCTTCACTGCCCCACT |
| sgRNA-E2-2# | GCGGCGCGCACAAACCTCCA |
| sgRNAs for dCas9 guided CAPTURE | |
| sgRNAs-E1 | E1-1: GTTTAGCAAAGGAATCAGAA  E1-2: GCAGTAAGGAGGGGGTAATG  E1-3: GGGGGGCAGTTAAGGCCGCT  E1-4: GAGCATCCCCCCTTTCCAAG  E1-5: GGAGCGTGATGTCATCAGCT  E1-6: GCTGTTTGGGGCGGAGAGGG  E1-7: GGAGGGTGGGGGCAGGCTGG  E1-8: GACAGTGAAAGACTGGAATG  E1-9: AGCCCCACCAAGAAGCTGGA  E1-10: AAGTCCAGGATGGGCAAGCA  E1-11: GGGAGAAACAGAGGTGAGGA  E1-12: AAAGGGGACCCACCAGGCGG |
| sgRNAs-E2 | E2-1: GAGACAGGATGGAGCGCCCT  E2-2: GCGGAGTAGTGGCGGCTGTC  E2-3: GGGTGGGGTTCGGGGATCGG  E2-4: GTGCGTGCCGACTCCTGCCC  E2-5: GAATTCCTGCGGGCGGAGCC  E2-6: GTGCGCGCCGCGCCCCCACC  E2-7: GGGGCACCGCCGTGAAAGGG  E2-8: TGCGCCTTCACTGCCCCACT  E2-9: GCAGGAGGCCTCGTAGGGAA  E2-10: GGACTCACGAGATACCAGGG  E2-11: GCGGCGCGCACAAACCTCCA  E2-12: TTAGCAGAAATAAGGAGCCC |

| **Table S3. Primers for RT-qPCR, related to STAR Methods.** | | | | |
| --- | --- | --- | --- | --- |
| Name | | Forward | | Reverse |
| YAP | | TAGCCCTGCGTAGCCAGTTA | | TCATGCTTAGTCCACTGTCTGT |
| TAZ | | GATCCTGCCGGAGTCTTTCTT | | CACGTCGTAGGACTGCTGG |
| CTGF | | AAAAGTGCATCCGTACTCCCA | | CCGTCGGTACATACTCCACAG |
| CRY61 | | CTCGCCTTAGTCGTCACCC | | CGCCGAAGTTGCATTCCAG |
| AJUBA | | ATGGGGAAGTCCTATCATCCAG | | TGGTAGTCGGTGACACAGTAT |
| seRNA | | TTACCCTGACTTCCGGAGGA | | GCTCCCTAGCATACGCGGAG |
| CDC6 | | CCAGGCACAGGCTACAATCAG | | AACAGGTTACGGTTTGGACATT |
| KIF23 | | AGTCAGCGAGAGCTAAGACAC | | GGTTGAGTCTGTAGCCCTCAG |
| RCC1 | | CGGCCCTGGTATCCATTCC | | CACTTTTGCTTAGACACACGGT |
| SGOL4 | | AACTCAGCAGTCACCTCATCT | | TGCACCTACGTTTAGGCAGAG |
| CENPF | | CTCTCCCGTCAACAGCGTTC | | GTTGTGCATATTCTTGGCTTGC |
| CDKN3 | | TCCGGGGCAATACAGACCAT | | GCAGCTAATTTGTCCCGAAACTC |
| KIF2C | | CTGTTTCCCGGTCTCGCTATC | | AGAAGCTGTAAGAGTTCTGGGT |
| MSH2 | | AGGCATCCAAGGAGAATGATTG | | GGAATCCACATACCCAACTCCAA |
| POC1A | | GACTCATGCCTCATGGTCTGG | | AGATGCGGACAGTCTTGTCTC |
| MELK | | TCTCCCAGTAGCATTCTGCTT | | TGATCCAGGGATGGTTCAATAGA |
| KNTC1 | | ACCTGAGTGTCGGTTCAAGAA | | CACTGATTGGTCGGCTACAATAA |
| CDK1 | | AAACTACAGGTCAAGTGGTAGCC | | TCCTGCATAAGCACATCCTGA |
| CDC7 | | GAGGCGTCTTTGGGGATTCAG | | GGTCCTACTTGTAACTGTGCTG |
| E2F3 | | AGAAAGCGGTCATCAGTACCT | | TGGACTTCGTAGTGCAGCTCT |
| GAPDH | | GGAGCGAGATCCCTCCAAAAT | | GGCTGTTGTCATACTTCTCATGG |
| **Table S4. Primers for CHIP, CHIP-qPCR,** **Enhancer KO validation and seRNA RACE related to STAR Methods.** | | | | |
| Name | Forward | | Reverse | |
| E1 | GAGGTGGCTGAACTCTGACC | | GGCGACAGTGAAAGACTGGA | |
| E2 | ACTTGGTCCCTTCCCTACGA | | CCCCCTGGTATCTCGTGAGT | |
| Promoter | GCTAGCCACAGACCGAACTT | | CAGAACATGCCTCCTGTCGT | |
| E1KO | CCCAggTgTCTACATTATCAgggATAgg | | ACACTTTTCTCTTCTCAggCACCT | |
| E2KO | ATCAATACTTCCTgAACAgAgTAggggAg | | CgCggAgAACCCTCgg | |
| Gene specific primers for seRNA RACE | | | | |
|  | 3' GSP primer | | GCAGGAAGGGGACAGTGCTCTGGGGC | |
|  | 5' GSP primer | | GGGTCCGAGGTGAAGACGCGGG | |

| **Table S5. Primers for site-directed mutagenesis, related to STAR Methods.** | | |
| --- | --- | --- |
| Name | Forward | Reverse |
| E1-MUT | CCTCCCCACCGCACAGTCTTTCACTGTCGCCCCAGGC | TGAAAGACTGTGCGGTGGGGAGGGTGGGGGCAGGCT |
| E2-MUT | CATTCTCCGACGCACTGCGGGCGGAGCCTGGGCCCGGG | CCGCCCGCAGTGCGTCGGAGAATGCGCGGGTGGGGG |
| Promoter-MUT | GGCATGGTGTGCGGTGTCGCGTCCCGGGACCGGACG | GACGCGACACCGCACACCATGCCCCTCCGCGCATTCCAA |
| YAP-4LE-MUT-1 | cgagagcagcaagagcagATGGAGAAGGAGAGGCTGCG | ctgctcttgctgctctcgCATCTGTTGCTGCTGGTTGGA |
| YAP-4LE-MUT-2 | cggctgaaacagcaagaagagCTTCGGCAGGCAATGCGG | ttcttgctgtttcagccgctcCCTCTCCTTCTCCATCTGCTCTT |
